# A Comprehensive Evaluation of Methods for Mendelian Randomization Using Realistic Simulations and an Analysis of 38 Biomarkers for Risk of Type-2 Diabetes

**DOI:** 10.1101/702787

**Authors:** Guanghao Qi, Nilanjan Chatterjee

## Abstract

**Background:** Mendelian randomization (MR) has provided major opportunities for understanding the causal relationship among complex traits. Previous studies have often evaluated MR methods based on simulations that do not adequately reflect the data-generating mechanism in GWAS and there are often discrepancies in performance of MR methods in simulations and real datasets.

**Methods:** We use a simulation framework that generates data on full GWAS for two traits under realistic model for effect-size distribution coherent with heritability, co-heritability and polygenicity typically observed for complex traits. We further use recent data generated from GWAS of 38 biomarkers in the UK Biobank to investigate their causal effects on risk of type-2 diabetes using externally available GWAS summary-statistics.

**Results:** Simulation studies show that weighted mode and MRMix are the only two methods which maintain correct type-I error rate in a diverse set of scenarios. Between the two methods, MRMix tends to be more powerful for larger GWAS while the opposite being true for smaller sample sizes. Among the other methods, random-effect IVW, MR-Robust and MR-RAPS tend to perform best in maintaining low mean squared error when the InSIDE assumption is satisfied, but can produce large bias when InSIDE is violated. In real data analysis, some biomarkers showed major heterogeneity in estimates of their causal effects on risk of type-2 diabetes across the different methods, with patterns similar to those observed in simulation studies.

**Conclusions:** Relative performance of different MR methods depends heavily on sample sizes of underlying GWAS, proportion of valid instruments and validity of the InSIDE assumption.

**Key Messages:**

- Many previous simulations studies to evaluate Mendelian randomization methods do not adequately reflect the data-generating mechanism of genome-wide association studies (GWAS).
- We use a simulation framework that generates data on full GWASs under realistic model informed by recent studies on effect-size distribution. We also used very recent GWAS data available on a large number of biomarkers to evaluate their causal effect on type-2 diabetes using alternative methods.
- Among the 10 methods that were compared, relative performance of different methods depends heavily on sample sizes of underlying GWAS, proportion of valid instruments and validity of the InSIDE assumption.
- Weighted mode and MRMix are the only two methods that maintain correct type I error rate in a diverse set of scenarios.

## Introduction

Epidemiological associations are often biased by unobserved confounders. Mendelian randomization (MR) – a form of instrumental variable approach that uses genetic variants as instruments – has provided major opportunities for understanding the causal relationship across complex traits [1–3]. The validity of early MR methods relied on a crucial assumption that the genetic variants have no effects on the outcome that are not mediated by the exposure. This assumption can be violated in the presence of “horizontal pleiotropy”. Recent studies have found that pleiotropy is a wide-spread phenomenon [4–9], leading to concerns over the accuracy of Mendelian randomization analysis. To deal with this challenge, many methods have been proposed that take advantage of the multitude of genetic instruments to reduce the bias due to horizontal pleiotropy. Different methods deal with different kinds of pleiotropy and often rely on different assumptions [7, 10–14].

Choosing a method for MR analysis can be challenging. While a number of previous studies have conducted various simulation studies to evaluate MR methods under alternative modeling assumptions, conclusions may be limited because these studies often do not incorporate realistic model for genetic architecture of complex traits as implied by recent studies of heritability/co-heritability [4, 15–19] and effect-size distribution [20–23]. Further, many simulation studies also directly simulate data on the instruments ignoring the process that instruments in reality are selected to be SNPs that reach genome-wide significance in an underlying genome-wide association study - as a result of which there should be a close relationship between sample size, number of available instruments, their average effect-sizes and precision of their estimated effects. Previous MR studies have used a fixed number of IVs and a fixed sample size [7, 14, 24] or vary one of them without the other [10–12, 25, 26], and generate the effects of genetic instruments from a fixed distribution without varying with the sample size or the number of IVs. In addition, genetic effects on exposure and outcomes are often simulated using uniform distributions while clearly many studies have shown they more likely to follow a spike-and-slab type distributions [22, 23, 27]. The magnitude of genetic effects simulated is also unrealistic in some studies such as only 25 SNPs explaining as much as 94% of variance of the exposure [10] or selected IVs explaining larger variance of the outcome than the exposure [28]. Because of these issues, performance of MR methods, in absolute and relative terms, can be discrepant between simulation studies and real GWAS datasets.

In this paper, we use a simulation framework that closely mirror real genome-wide association studies. In particular, we simulate data on genome-wide set of SNPs and select instruments based on SNPs which reach genome-wide significance in the underlying study of the exposure. We simulate genetic effects constrained by realistic values for heritability/co-heritability and models for effect-size distribution [4, 15–23]. We compare performance of a variety of existing methods under different sample sizes of underlying GWAS and correspondingly number of IVs and vary the proportion of valid instruments and mechanisms of pleiotropy. We further compare the methods in real data analysis to estimate the effect of blood and urine biomarkers on type 2 diabetes. Results from these simulation studies provide comprehensive and realistic insights into strengths and limitations of existing methods.

## Methods

We begin by introducing a few notations. Let *X* denote the exposure, *Y* denote the outcome and *U* denote a potential confounder. Let *G*_*j*_ denote the genotype of SNP *j*. Throughout most MR literature [10–12, 26, 28], the simulations are conducted using the following model

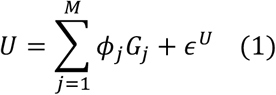

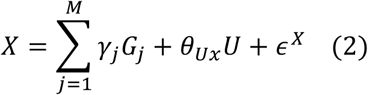

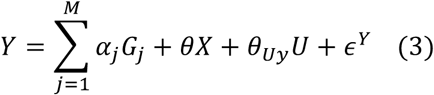

Here *ϕ*_*j*_, *γ*_*j*_, *α*_*j*_ denote the direct effect of SNP *j* on *U, X* and *Y*, respectively. We also adopt this model in our simulations. But unlike most previous studies that simulate only the selected instruments, we generate data for all common variants in the genome. We are interested in estimating the causal effect (*θ*) of *X* on *Y*. The effects of the confounder *U* on *X* and *Y* are denoted by *θ*_*Ux*_ and *θ*_*Uy*_, respectively. The error terms *ϵ*^*U*^, *ϵ*^*X*^ and *ϵ*^*Y*^ are independent and normally distributed with mean 0.

We generated data from model (1)-(3) using 200,000 independent SNPs as representative of all underlying common variants. We generate *ϕ*_*j*_, *γ*_*j*_, *α*_*j*_ from mixture normal distributions which have been shown to be appropriate for modeling effect-size distribution for complex traits in GWAS [20–23]. Under the above model, when the confounder *U* has heritable component, the InSIDE assumption [10] is violated as direct and indirect effect of some SNPs on the outcome are correlated due to mediation by common factor *U*.

### Balanced Horizontal Pleiotropy with InSIDE Assumption Satisfied

We first simulate settings where SNPs with direct effect on *X* can also have direct effect on *Y*, thus allowing horizontal pleiotropy, but we allow the InSIDE assumption to be satisfied by setting *ϕ*_*j*_ = 0 for all SNPs. We generate *γ*_*j*_ and *α*_*j*_ across SNPs from the following distribution:

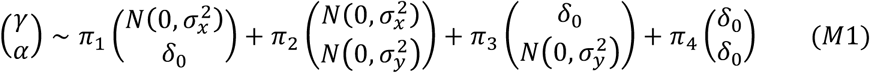

In the above, the first component is the set of valid IVs, i.e. the SNPs which have no horizontal pleiotropic effect on *Y* (*δ* _0_ is the point mass at 0). The second component is the set of SNPs with horizontal pleiotropic effect on *Y*. However, because here we assume *γ*_*j*_ and *α*_*j*_ are independent, in this setting the InSIDE assumption is satisfied. The third component is the set of SNPs that are only associated with *Y* and the fourth component is the set of SNPs that have no association with either trait. Mixture proportions are denoted by *π*_1_, *π*_2_, *π*_3_, *π*_4_ and variance parameters are denoted by 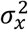 and 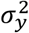 (see **Table 1** and **Supplementary Table 1** for details on values used for the parameters).

**Table 1.** Choice of parameters in simulation studies where the true causal effect of *X* on *Y* is 0.2 and 50% of the potential IVs are valid.

| Setting | # of causal variants | Heritability of $X$ | Heritability of $Y$<sup>a</sup> | Genetic correlation<sup>b</sup><br>due to<br>vertical/horizontal<br>pleiotropy |
| --- | --- | --- | --- | --- |
| Balanced pleiotropy,<br>InSIDE satisfied<br>[Model (M1), Methods] | 200,000 independent SNPs<br>representing the common<br>variants in the genome.<br>1) 2,000 SNPs have direct effect<br>on $X$ but not $Y$ (addressed as<br>**potential valid IVs**).<br>2) 2,000 SNPs have direct effects<br>on both $X$ and $Y$ (addressed as<br>**SNPs in horizontal pleiotropy**).<br>3) 2,000 SNPs have direct effects<br>on $Y$ but not $X$.<br>4) The remaining 194,000 SNPs<br>have no effects on either $X$ or $Y$. | **Total 20%:** 10% due to potential valid IVs;<br>10% due to SNPs in horizontal pleiotropy. | **Total 20.8%:** 0.8% due to causal effect of $X$;<br>10% due to SNPs in horizontal pleiotropy;<br>10% due to SNPs that have effects on $Y$ but not $X$. | 0.196 / 0 |
| Balanced pleiotropy,<br>InSIDE violated<br>[Model (M2), Methods] | | **Total 20%:** 10% due to potential valid IVs;<br>8.2% due to direct effects of SNPs in<br>horizontal pleiotropy; 1.8% due to<br>confounder $U$. | **Total 21.52%:** 0.8% due to causal effect of $X$;<br>2.52% due to confounder $U$ and its covariance with $X$;<br>8.2% due to direct effects from SNPs in horizontal<br>pleiotropy that are not mediated by $U$;<br>10% due to SNPs that have effects only on $Y$. | 0.193 / 0.087 |
| Directional pleiotropy,<br>InSIDE satisfied<br>[Model (M3), Methods] | | **Total 20%:** 10% due to potential valid IVs;<br>10% due to SNPs in horizontal pleiotropy. | **Total 30.8%:** 0.8% due to causal effect of $X$;<br>15% due to SNPs in horizontal pleiotropy (10%<br>balanced + 5% directional effect);<br>15% due to SNPs that have effects only on $Y$. | 0.161 / 0 |
| Directional pleiotropy,<br>InSIDE violated<br>[Model (M4), Methods] | | **Total 20%:** 10% due to potential valid IVs;<br>8.2% due to direct effects of SNPs in<br>horizontal pleiotropy; 1.8% due to<br>confounder $U$. | **Total 31.52%:** 0.8% due to causal effect of $X$;<br>2.52% due to confounder $U$ and its covariance with $X$;<br>13.2% due to direct effects from SNPs in horizontal<br>pleiotropy that are not mediated by $U$ (8.2% balanced<br>+ 5% directional effect);<br>15% due to SNPs that have effects only on $Y$. | 0.159 / 0.072 |

See Supplementary Table 1 for the exact choice of parameters in all settings.
<sup>a</sup> When directional pleiotropy exists, the direct SNP effects on $Y$ follow distribution $N(\mu_y, \tilde{\sigma}_y^2)$ , which can be decomposed into $N(0, \tilde{\sigma}_y^2) + \mu_y$ . “Balanced effects” refer those from the first component $N(0, \tilde{\sigma}_y^2)$ and “directional effects” refer to those from the second component $\mu_y$ .
<sup>b</sup> Genetic correlation between two traits $X$ and $Y$ are defined as $h_{xy}/\sqrt{h_x^2 h_y^2}$ , where $h_{xy}$ is the covariance of the genetic components of $X$ and $Y$ , $h_x^2$ and $h_y^2$ are the heritability of $X$ and $Y$ respectively.

### Balanced Horizontal Pleiotropy with InSIDE Assumption violated

Next we allow the InSIDE assumption to be violated by allowing a fraction of SNPs to have an effect on the confounder *U*. Here we generate *ϕ*_*j*_, *γ*_*j*_, *α*_*j*_ from the tri-variate normal mixture:

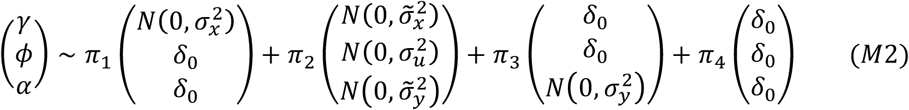

In the above, the first component corresponds to valid instruments which have only direct effect on *X*. The second component allows a set of SNPs that have effects on *U* and thus creating horizontal pleiotropic effect with the InSIDE assumption violated. We also allow the same set of SNPs to have direct effects on *X* and *Y*, but the effect sizes are of smaller magnitude.

### Directional Pleiotropy

In the above two settings, we assumed direct effects of the SNPs on the outcome *Y* have mean zero. Next to simulate directional pleiotropy, we generate *α*_*j*_ from a distribution with non-zero mean (*μ*_*y*_). When the InSIDE assumption holds, we simulate from

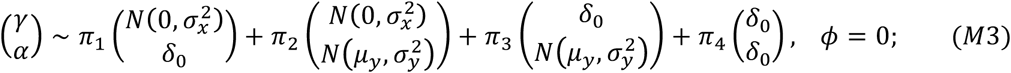

when InSIDE does not hold, we simulate from

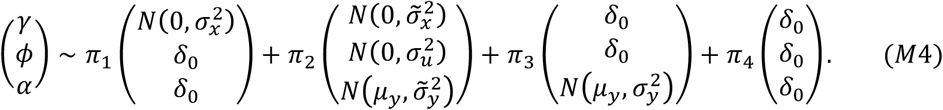

### Simulating Data on Genome-wide Association Studies

We simulate individual level data for independent genome-wide association studies for *X* and *Y* following the above model when sample size is not too large (*N* ≤ 100*k*). However, for very large sample size, generation and analysis of individual level data can become computationally prohibitive and we simulate summary-level association statistics directly (addressed as *summary-level simulations*). We observe that the total effects of SNPs on *X* and *Y* are implied by model (1)-(3):

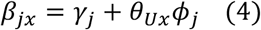

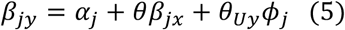

Thus, we simulate *γ*_*j*_, *ϕ*_*j*_ and *α*_*j*_ s as before and then directly generate summary statistics by adding to *β*_*jx*_ and *β*_*jy*_ error terms with mean 0 and variance inversely proportional to the sample size. Details can be found in the **Supplementary Note 1**.

### Choosing Parameters Values Reflecting Realistic Genetic Architecture

We found previous studies for evaluation of MR methods have often not followed realistic model for genetic architecture of complex traits. In particular, a very recent study that evaluated a large number of methods for MR analysis following the basic setup described in (1)-(3), used highly unrealistic parameter settings [28]. The study assumes the instruments explain an unrealistically large proportion of the variance of *X* and *Y* and ignore the relationship between sample size, number of instruments and effect-size distributions (**Supplementary Table 2, Supplementary Note 2**).

We chose parameter values in our model so that they reflect realistic genetic architecture of complex traits and results from our simulated GWAS track what are typically observed in empirical studies. In **Table 1**, we show the parameter values chosen for different simulation settings and corresponding values of heritability of the two traits and their co-heritability due to horizontal and vertical pleiotropy. Further in **Figure 1**, we show how under simulation settings as the sample size for GWAS of *X* increases, the number of available IVs and the amount of variance they explain for the two traits increase. These patterns closely correspond to that observed in GWAS of many traits, such as BMI. Further see **Supplementary Table 1** for the exact choice of parameters.

**Figure 1.**
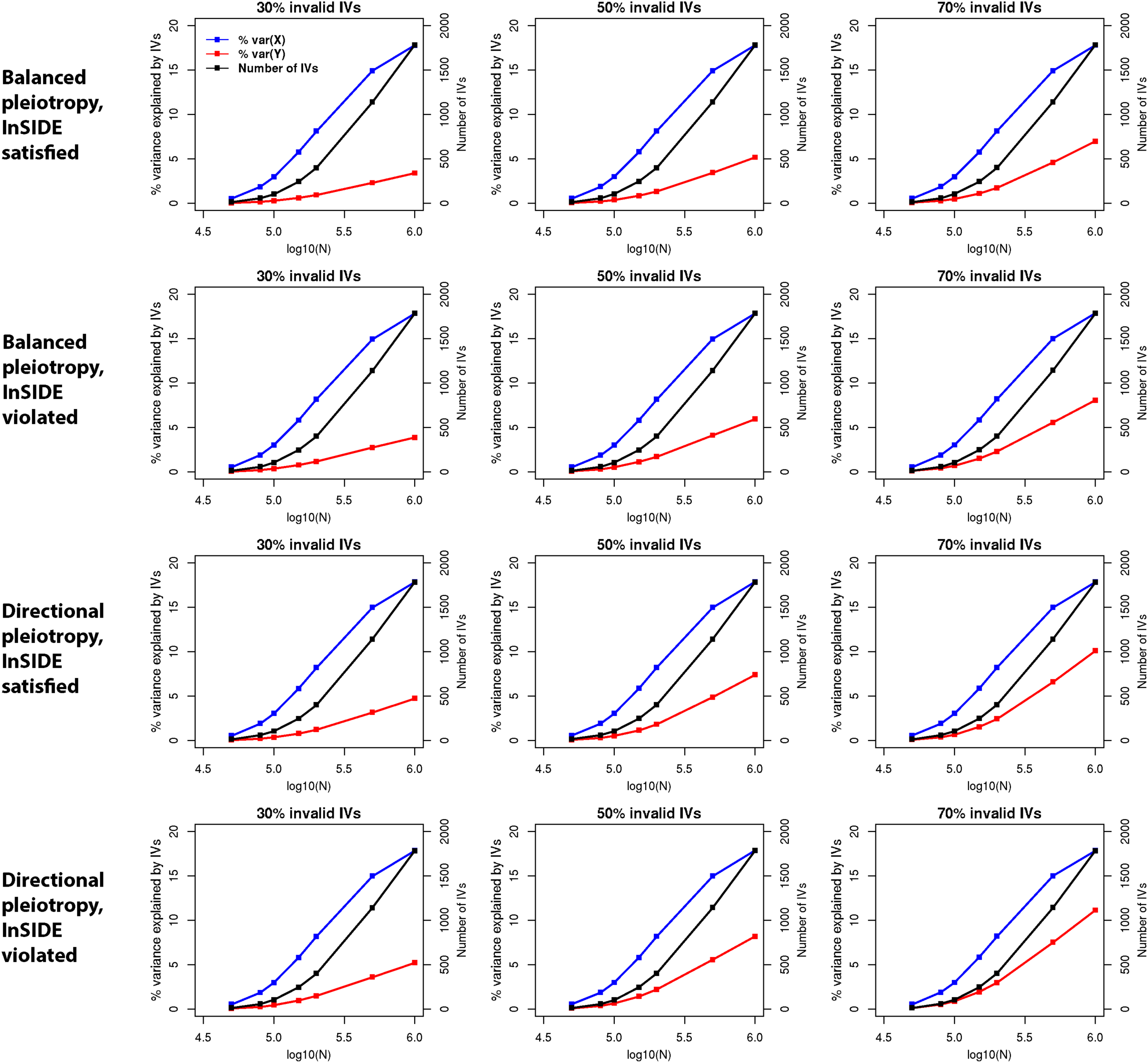
Relationship among sample size, average number of IVs and variance of traits explained by the IVs under different scenarios of simulation studies. The true causal effect from *X* to *Y* is 0.2. Sample size of the study associated with *X* is *N*; sample size of the study associated with *Y* is *N*/2. IVs are defined as the SNPs which reach genome-wide significance (z-test *p* < 5×10^−8^) in the study associated with *X*. Averages are calculated over 200 simulations.

### Existing Robust MR Methods

We compare all nine methods investigated in the recent study indicated above [28], which include weighted median [11], weighted mode [12], MR-Egger [10], MR-Robust, MR-Lasso [25], MR-RAPS [29], MR-PRESSO [7], MRMix [13] and contamination mixture (Con-mix) [14]. In addition, we include the inverse-variance weighted method with multiplicative random effects (IVW-r) in comparison [30]. See **Supplementary Note 3** for a summary of the different methods and **Supplementary Table 3** for the software and tuning parameters used to implement the methods.

### MR Analysis for Biomarker effects on Type 2 diabetes

We applied the variety of available methods for MR analysis to investigate causal effect of 38 blood and urine biomarker measures in the UK Biobank study [31] on the risk of type-2 diabetes (T2D). We accessed the summary statistics from recent analysis of the biomarkers in the UK Biobank (N=318 984) [31]. We restricted the analysis to those biomarkers which have more than 25 associated instruments and the instruments explain at least 1% of the biomarker variance. On the outcome side, we accessed summary statistics from the largest GWAS on type 2 diabetes which consists of 74 124 T2D cases and 824 006 controls [32]. Details are included in **Supplementary Note 4**.

## Results

We present main results under the simulation scheme that generate summary-level data directly as it allows exploration of GWAS of very large sample size (*N* > 100*k*). Under smaller sample size where we did simulation with both individual and summary-level data, we see the results are very comparable across the two schemes (see **Supplementary Figures 1 and 2**).

**Figure 2.**
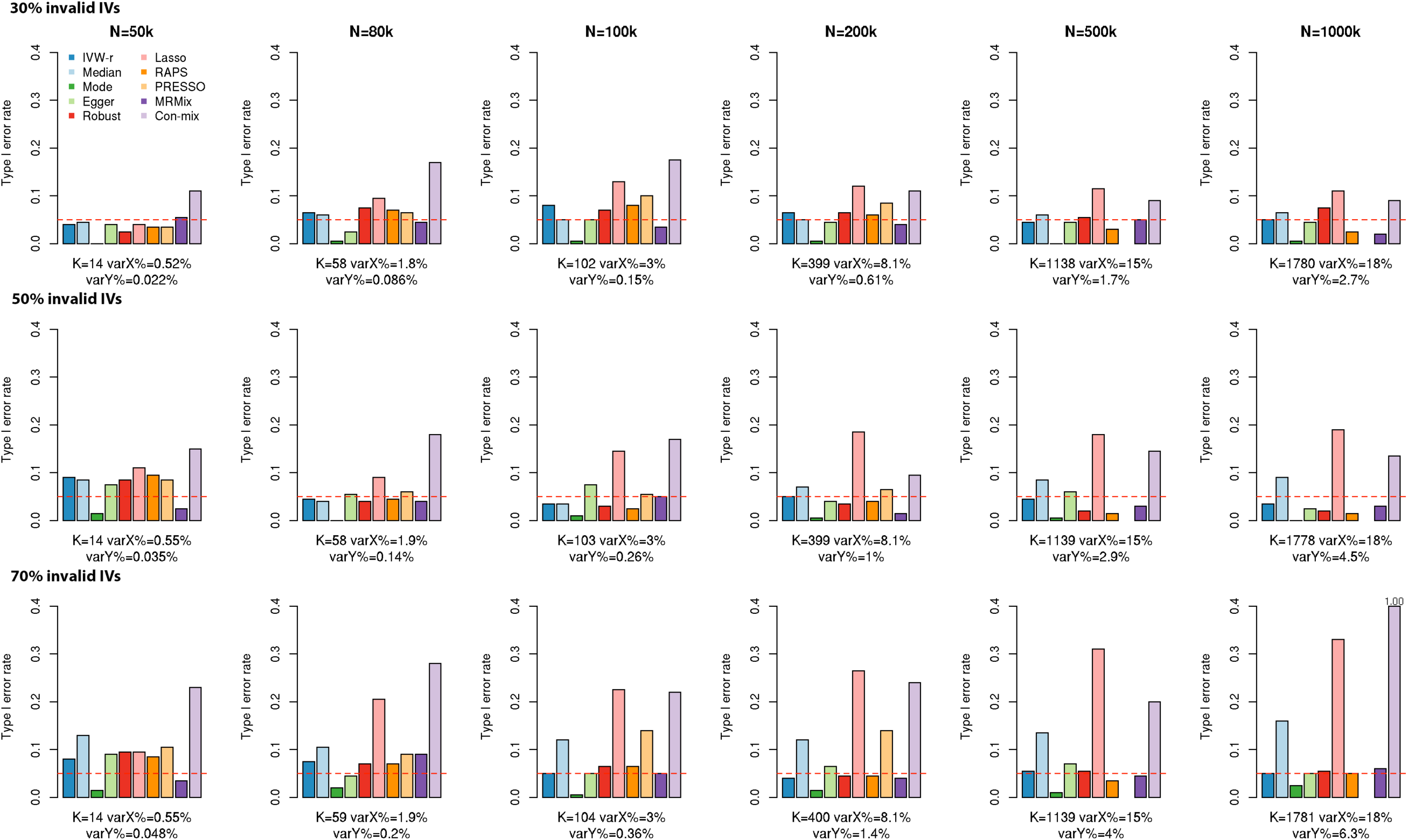
Type I error rates for alternative MR methods in simulations with balanced pleiotropy and InSIDE assumption satisfied. There is no true causal effect from *X* to *Y*. Empirical type I error rates are reported over 200 simulations. Sample size of the study associated with *X* is *N*; sample size of the study associated with *Y* is *N*/2. *K*: the average number of IVs, defined as the SNPs which reach genome-wide significance (z-test *p* < 5×10^−8^) in the study associated with *X*; varX%: average percentage of variance of *X* explained by IVs; varY%: average percentage of variance of *Y* explained by IVs. The red dashed line is the nominal significance threshold 0.05.

Under balanced pleiotropy and InSIDE assumption, the weighted mode estimator and MRMix controls type I error at the nominal level across different scenarios (**Figure 2**). The type I error rate of weighted mode usually falls far below the nominal value while that of MRMix generally remains fairly close to the nominal value. Among other methods, IVW-r, MR-Egger, MR-Robust and MR-RAPS are the most robust as they generally maintain the nominal type I error except when the number of invalid IVs are large or/and sample size is small. The least robust methods are MR-Lasso and Con-mix which often have extremely high type I error reaching even up to 100% for the latter method when *N* = 1000*k* and 70% of the instruments were invalid. When we compare the methods in terms of MSE (**Figure 3**), we find that IVW-r, weighted median, MR-Robust, MR-Lasso, MR-PRESSO and MR-RAPS perform comparably to each other and have an advantage over the others. Among weighted mode and MRMix, which are the only two methods maintaining nominal type-I error, we find MSE for weighted mode could be much smaller than MRMix for smaller sample size (e.g *N* = 50*k*) but the latter method has a clear advantage when sample size becomes larger (*N* ≥ 200*k*). Power tracks closely with the MSE among the methods that have reasonably well controlled or only moderately inflated type I error rates (**Supplementary Figure 3**).

**Figure 3.**
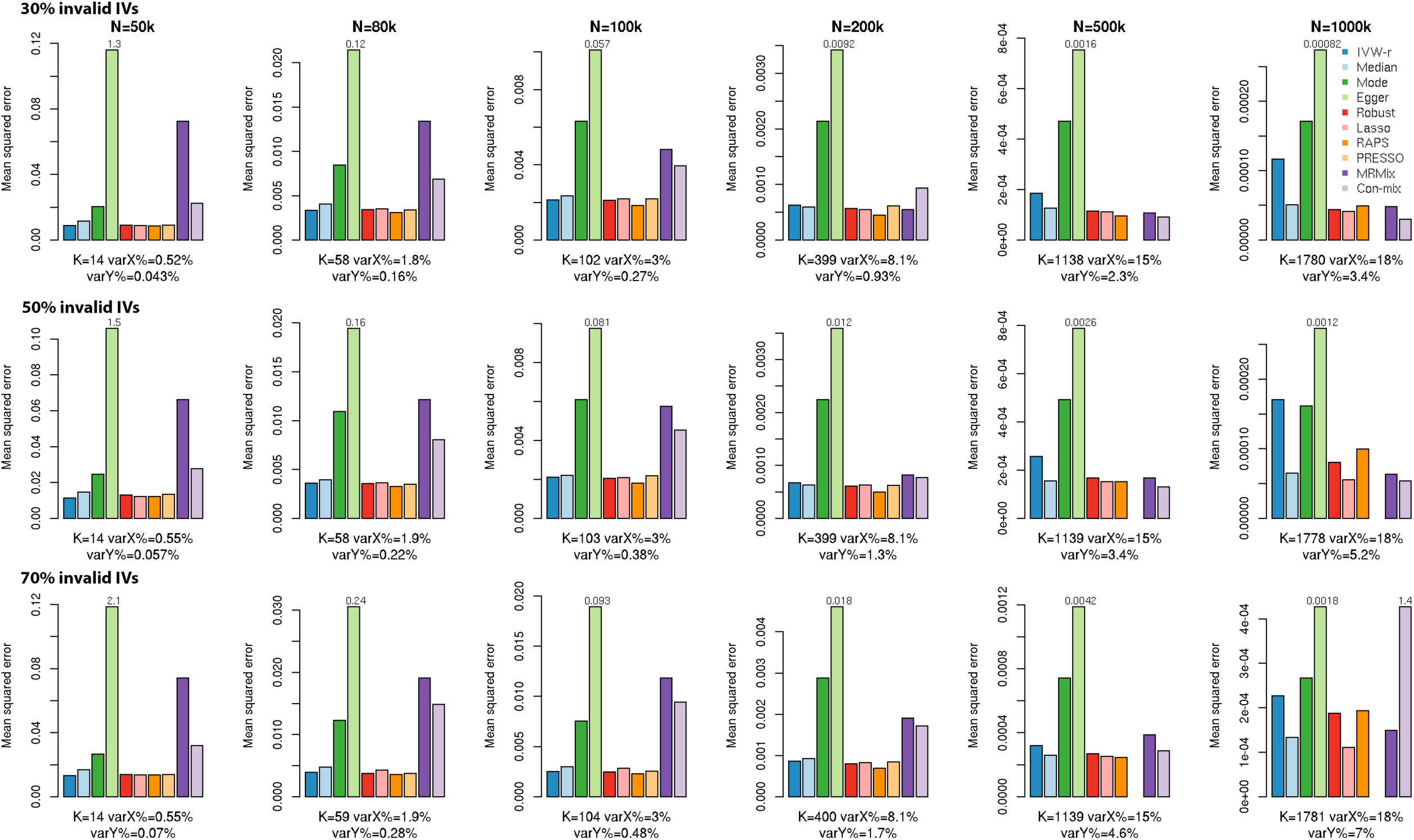
Mean squared error for estimation of causal effect under alternative MR methods in simulations with balanced pleiotropy and InSIDE assumption satisfied. The true causal effect from *X* to *Y* is 0.2. Mean squared errors are reported over 200 simulations. Sample size of the study associated with *X* is *N*; sample size of the study associated with *Y* is *N*/2. *K*: the average number of IVs, defined as the SNPs which reach genome-wide significance (z-test *p* < 5×10^−8^) in the study associated with *X*; varX%: average percentage of variance of *X* explained by IVs; varY%: average percentage of variance of *Y* explained by IVs. Bars higher than the upper limit of the panel are truncated and marked with the true value.

We also compared the different methods in terms of bias and variance separately. Under balanced pleiotropy and InSIDE assumptions, all MR methods generally show some bias when the sample size is small and gradually the bias diminishes as the sample size increases (**Supplementary Figure 4**). When there is no causal effect, all methods give average estimates of causal effect close to 0 across wide range of sample sizes except for Egger regression with *N* ≤ 100*k* and Con-mix with *N* = 1000*k* and large number of invalid IVs. Comparing the empirical versus estimated standard errors of the methods we observe that the MR-Lasso and Con-mix produce severely underestimated standard errors (**Supplementary Figure 5**), while weighted median and MR-PRESSO has underestimated standard error when 70% of the instruments are valid. This is likely to be the reason for type I error inflation. Throughout the settings IVW-r, MR-Egger, MR-Robust and MR-RAPS give accurate standard error estimation; weighted mode has overestimated standard error which is likely to be the reason for its overly conservative type I error rate. The standard error estimate of MRMix tends to be too large when *N* ≤ 100*k*, but converges to the truth when *N* ≥ 200*k*.

When the InSIDE assumption is violated, we find all methods except weighted mode and MRMix could have extremely inflated type I error (**Figure 4**). As before while MRMix maintains type I error close to the nominal level, weighted mode is very conservative. Among other methods, Con-mix and MR-Egger are less biased, but even these methods have unacceptably high type I error in a variety of scenarios. When the methods are compared in terms of MSE (**Figure 5**), for smaller sample sizes (*N* ≤ 100*k*), the methods IVW-r, weighted median, MR-Robust, MR-Lasso and MR-PRESSO seem to be the best even though they may have appreciable bias. For larger sample size (*N* ≥ 200*k*), MRMix generally has the smallest MSE; and had substantially higher power than weighted mode, which also controls the type I error (**Supplementary Figure 6**). When we inspected bias and variance separately, it is evident that when the InSIDE assumption is violated all methods, except weighted mode and MRMix, can have large bias, even when there is no causal effect, and this bias does not disappear with increasing sample size (**Supplementary Figure 7**). The patterns of bias in standard error estimation for the different methods are similar as what we described before when the InSIDE assumption is satisfied (**Supplementary Figure 8**).

**Figure 4.**
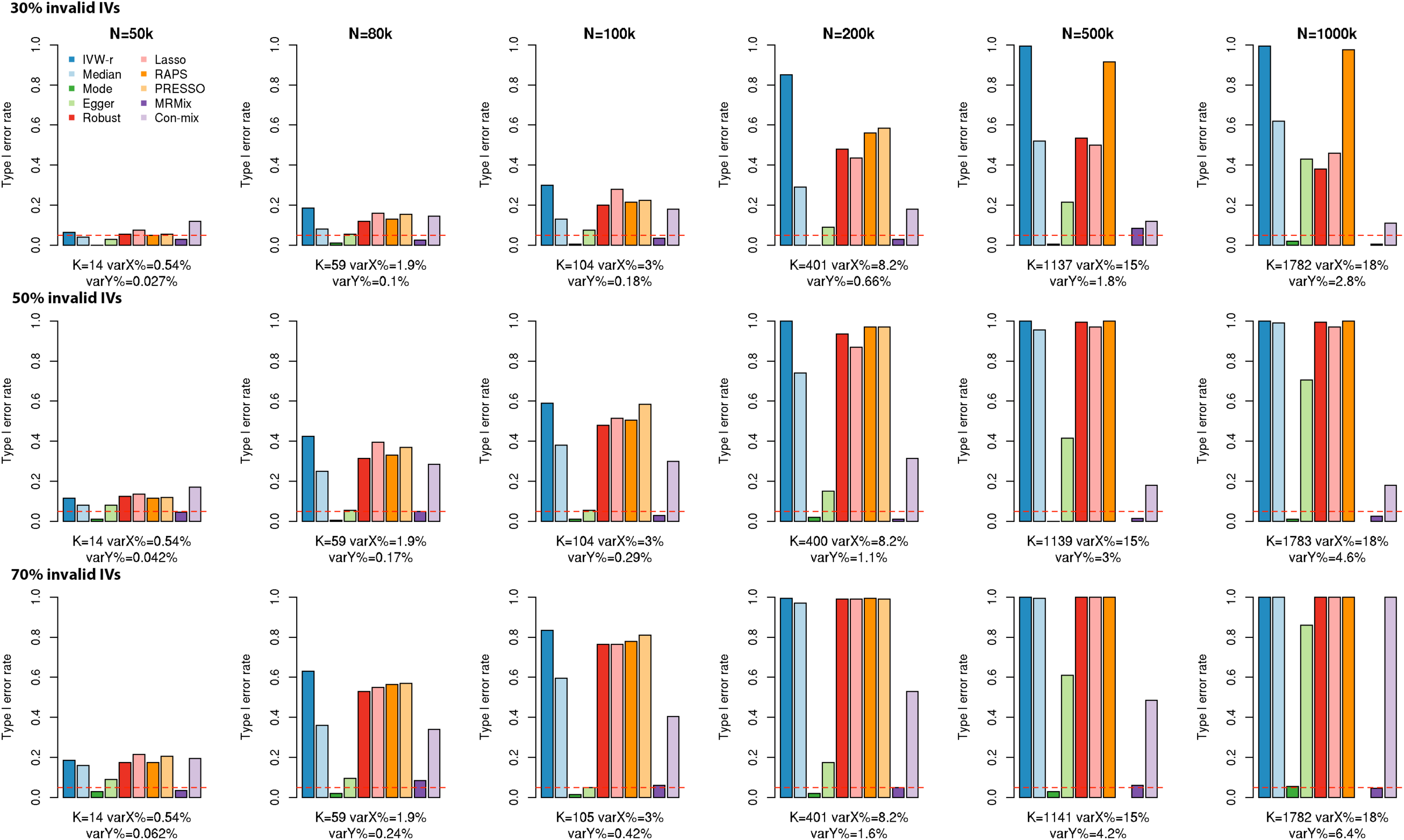
Type I error rate in in simulations with balanced pleiotropy and InSIDE assumption violated. There is no true causal effect from *X* to *Y*. Type I error rates are reported over 200 simulations. Sample size of the study associated with *X* is *N*; sample size of the study associated with *Y* is *N*/2. *K*: the average number of IVs, defined as the SNPs which reach genome-wide significance (z-test *p* < 5×10^−8^) in the study associated with *X*; varX%: average percentage of variance of *X* explained by IVs; varY%: average percentage of variance of *Y* explained by IVs. The red dashed line is the nominal significance threshold 0.05.

**Figure 5.**
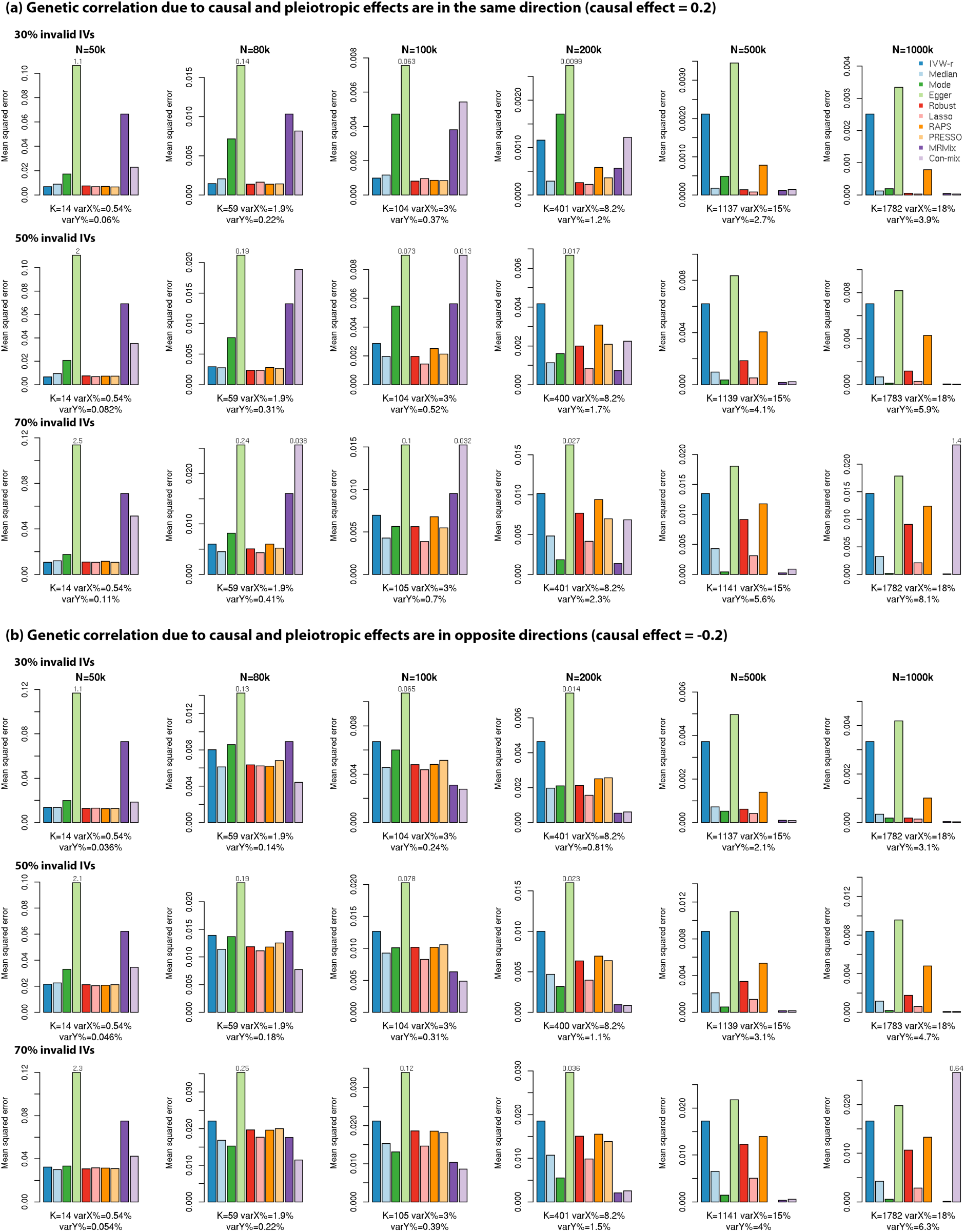
Mean squared error in in simulations with balanced pleiotropy and InSIDE assumption violated. Mean squared errors are reported over 200 simulations. Sample size of the study associated with *X* is *N*; sample size of the study associated with *Y* is *N*/2. *K*: the average number of IVs, defined as the SNPs which reach genome-wide significance (z-test *p* < 5×10^−8^) in the study associated with *X*; varX%: average percentage of variance of *X* explained by IVs; varY%: average percentage of variance of *Y* explained by IVs. Bars higher than the upper limit of the panel are truncated and marked with the true value.

**Figure 6.**
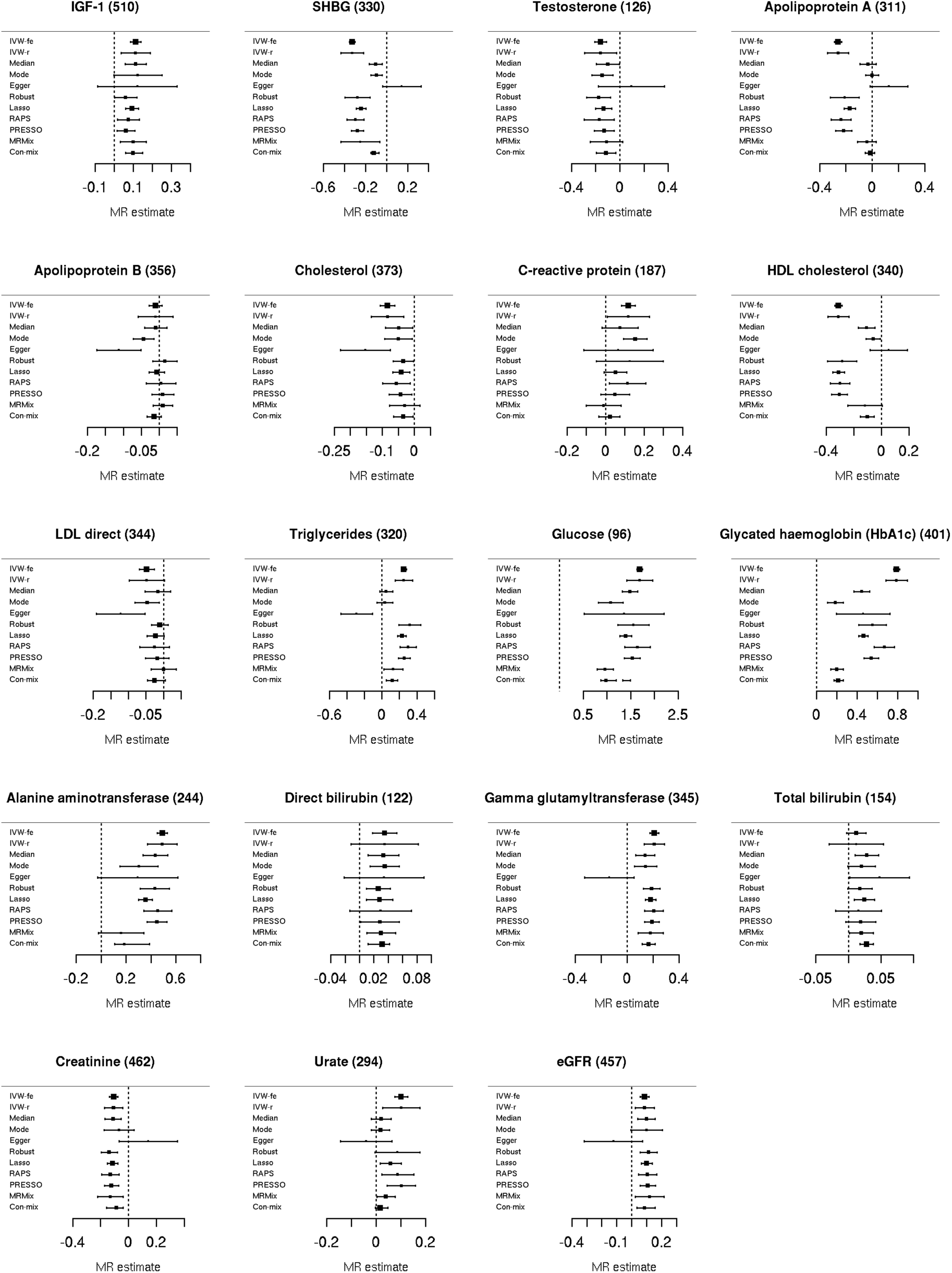
MR estimates and corresponding 95% confidence intervals of causal effects from UK Biobank blood and urine biomarkers to type 2 diabetes. MR estimate is the log-OR of type 2 diabetes (T2D) per SD increase in the exposure. We present the results for biomarkers that satisfy three criteria: 1) >25 Ivs 2) IVs explain >1% variance of the exposure 3) have a causal effect on T2D at *p* < 0.005 by at least one method (except IVW-fe). IVW-fe: IVW with fixed effects; IVW-r: IVW with random effects. Title of each panel is the name of the biomarker followed by number of IVs.

In the simulations with directional pleiotropy, the patterns are fairly similar with corresponding scenarios for balanced pleiotropy setting except that the type I error for a number of methods increased somewhat in certain settings (**Supplementary Figures 9-14**).

### Causal effects of biomarkers on type 2 diabetes

We further compared the methods in real data analysis to estimate the causal effects of 38 UK Biobank blood and urine biomarkers on type 2 diabetes. We focus on those 19 biomarkers which were found reported to have significant causal effect at *p* < 0.005 by at least one of the methods.

Unsurprisingly, all the methods consistently report that higher levels of glucose causally increase the risk of T2D (**Figure 6 and Supplementary Figure 15**). Another T2D biomarker, HbA1C, also has a substantial causal effect on T2D consistently across all the methods. However, substantial heterogeneity is observed in estimates of the causal effect across methods. Weighted mode, MRMix and Con-mix estimates the OR of T2D to be around 1.22 per SD unit increase in HbA1C; weighted median, MR-PRESSO, MR-Robust and MR-Lasso estimate this OR to be 1.57-1.73; IVW-r and MR-RAPS report a strong effect of OR=2.2 and 1.95, respectively. The next strongest causal effect on T2D was observed for alanine aminotransferase (ALT, a liver function biomarker) with some degree of heterogeneity in effect estimates across methods.

For ten biomarkers, estimates from all MR methods except Egger regression are consistent with each other (**Figure 6**). They include IGF-1, testosterone, apolipoprotein B, cholesterol, LDL direct, direct bilirubin, gamma glutamyltransferase (GGT), total bilirubin, creatinine and eGFR. Interestingly, these biomarkers generally have small to moderate effects on T2D. The results appear to be less homogeneous for the biomarkers that are shown to have a substantial effect on T2D (|log (*OR*)| > 0.25) by some methods (**Figure 6 and Supplementary Figure 15**). Consistent with what was observed in the simulation studies, IVW, MR-PRESSO, MR-Robust, MR-Lasso and MR-RAPS tend to give estimates that are similar to each other; weighted mode, MRMix and contamination mixture tend to give similar estimates, which are usually closer to the null than those by the other methods. For example, IVW, MR-PRESSO, MR-Robust, MR-Lasso and MR-RAPS report the OR of T2D to be ∼0.74 per SD unit increase in HDL cholesterol, and 1.26-1.38 per SD unit increase in triglycerides; in contrast, weighted mode, MRMix and contamination mixture report the OR to be 0.89-0.94 for HDL cholesterol and 1.03-1.14 for triglycerides. Similar patterns were also observed for glucose, HbA1C and ALT, which were discussed previously.

We also observed that the estimated confidence interval (CI) by Con-mix and MR-Lasso tend to be substantially smaller than those by the other methods. In a number of cases, e.g. total bilirubin and urate, Con-mix report confidence intervals of similar or smaller length to that by the fixed-effect IVW (IVW-fe). These results are consistent with simulation studies which indicated that the CI estimates by these two methods can be anti-conservative and can lead to type I error.

## Discussion

In this paper, we evaluate a variety of methods for polygenic MR analysis using a simulation framework which generate data closely resembling patterns observed in empirical genome-wide association studies. Results reveal varying performance of the MR methods under different scenarios. When the sample size is large (e.g. *n*_*X*_ > 200*k, n*_*y*_ > 100*k*), MRMix appears to be best or close to be best, whether or not InSIDE assumption is satisfied, in terms of its ability to control type I error rate and bias and yet maintaining relatively high power and low MSE. When the sample size is smaller (e.g. *n*_*X*_ ≤ 100*k, n*_*y*_ ≤ 50*k*), no method appears to be performing uniformly well across all scenarios. When the InSIDE assumption holds, IVW-r, MR-Robust and MR-RAPS lead to the smallest mean-squared errors and they usually have either well controlled or modestly inflated type I error rates. The weighted median method also performs well when the proportion of valid IVs is not too high (e.g. ≤ 30%) but suffers from more severe type I error when this proportion increases. When the InSIDE assumption is violated, only weighted mode and MRMix have well controlled type I error and all the other methods can have severely inflation in type I error. Between these two methods, weighted mode tends to be more efficient and powerful for moderate sample size (e.g. *n*_*X*_ ≤ 80*k, n*_*y*_ ≤ 40*k*).

We observe that the type I error of a number of methods are significantly affected due to not only bias in point estimation but also that of the underlying standard error estimators. In particular, we found that the type I error of the weighted mode can often be substantially lower than the desired nominal level due to conservativeness of underlying standard error estimator (**Supplementary Figure 5 and 8**). Further for a number of other estimators, which did not have bias in point estimation at least when the InSIDE assumption is satisfied, have inflated type I error due to anticonservative standard error estimation. It is possible that in the future the type I error or/and power for some of these procedures can be improved through implementation of more robust standard error estimation procedures.

Both similarities and differences exist between our simulation results and the results reported in a recent study also comparing most of the same MR methods [28]. Both studies found that the weighted mode estimator has well controlled type I error rate across various scenarios. The biggest difference is observed for mixture model based methods. In our study, MRMix is shown to perform well under large sample size, especially when the InSIDE assumption is violated. The study by Slob and Burgess restricted their simulation studies with sample size *n*_*X*_ = *n*_*y*_ = 10 000. Under such small sample size, MRMix can be unstable because the performance of the method depends on the ability of the underlying mixture model to cluster valid and invalid IVs based on underlying variance components. If the sample size is small and estimates of effect sizes for the IVs have large variability, then the two variance components are not well separable and the resulting estimates can have large uncertainty. When sample size increases, variance component associated with valid IVs are expected to be smaller than those with the non-valid IVs and then the method allows robust estimation of causal effects. In addition, we find the alternative mixture model-based method, Con-mix can have smaller MSE than that of MRMix specially for smaller sample sizes, but the former method can have much higher type I error across a variety of scenarios with or without the InSIDE assumption violated. We also observed a numerical breakdown of Con-mix for very large GWAS (e.g *n*_*X*_ = 1000*k*) for which the cause is not well understood.

The simulation framework we propose can be broadly useful for future evaluation of emerging MR methods. We simulate data on genome-wide panel of SNPs and apply a p-value threshold to select IVs as is done in real studies. This procedure naturally reflects the relationship among sample size, number of IVs and instrument strength. We also use realistic distributions to generate genetic effect sizes based on recent work on heritability and effect size distributions [4, 15–23]. We studied the performance of MR methods in a wide range of sample sizes and scenarios of violations of standard assumptions of MR analysis. We propose a framework for directly simulating summary-level data implied by the model for individual data for reducing the computational burden associated with simulating vary large GWAS.

Results from simulation studies and real data analysis were generally consistent. The number of instruments available for the different biomarkers and the associated variances explained were within reasonable range of scenarios considered in the simulation studies given the sample size. Both simulation studies and real data analysis show that IVW-r, MR-PRESSO, MR-Robust, MR-Lasso and MR-RAPS tend to give similar estimates of causal effects, while the estimates from weighted mode, MRMix and Con-mix are close to each other. Both simulation studies and real data analysis also indicated the estimated confidence intervals of MR-Lasso and Con-mix can be anti-conservative and thus lead to increased type-I error. (**Supplementary Figures 5 and 8, Figure 6**).

In summary, we conducted large-scale and realistic simulation studies to compare 10 methods for Mendelian randomization analysis. Our results show that while for GWAS with very large size the mixture model based method MRMix emerges as the most robust method, for medium to smaller sample sized studies there is no single method that performs uniformly well across all scenarios. Thus in real data analysis it is prudent to apply a few alternative methods with complimentary features and strengths and assess sensitivity of findings across all these methods.

## Code availability

The code for the simulation studies is available on GitHub: https://github.com/gqi/MR_comparison_simulations

## Acknowledgement

The work was supported by a RO1 grant from the National Human Genome Research Institute [1 R01 HG010480-01].

## References

1. Davey Smith G, Ebrahim S. ‘Mendelian randomization’: can genetic epidemiology contribute to understanding environmental determinants of disease. Int J Epidemiol. 2003;32:1–22.

2. Davey Smith G, Hemani G. Mendelian randomization: genetic anchors for causal inference in epidemiological studies. Hum Mol Genet. 2014;23:R89–R98.

3. Zheng J, Baird D, Borges M-C et al. Recent developments in Mendelian randomization studies. Current Epidemiology Reports. 2017;4:330–345.

4. Bulik-Sullivan B, Finucane HK, Anttila V et al. An atlas of genetic correlations across human diseases and traits. Nat Genet. 2015;47:1236–1236.

5. Pickrell JK, Berisa T, Liu JZ, Ségurel L, Tung JY, Hinds D. Detection and interpretation of shared genetic influences on 42 human traits. Nat Genet. 2016;48:709.

6. Sivakumaran S, Agakov F, Theodoratou E et al. Abundant pleiotropy in human complex diseases and traits. Am J Hum Genet. 2011;89:607–618.

7. Verbanck M, Chen C-Y, Neale B, Do R. Detection of widespread horizontal pleiotropy in causal relationships inferred from Mendelian randomization between complex traits and diseases. Nat Genet. 2018;50:693.

8. Visscher PM, Yang J. A plethora of pleiotropy across complex traits. Nat Genet. 2016;48:707–708.

9. Visscher PM, Wray NR, Zhang Q et al. 10 years of GWAS discovery: biology, function, and translation. The American Journal of Human Genetics. 2017;101:5–22.

10. Bowden J, Davey Smith G, Burgess S. Mendelian randomization with invalid instruments: effect estimation and bias detection through Egger regression. Int J Epidemiol. 2015;44:512–525.

11. Bowden J, Davey SG, Haycock PC, Burgess S. Consistent estimation in Mendelian randomization with some invalid instruments using a weighted median estimator. Genet Epidemiol. 2016;40:304–314.

12. Hartwig FP, Davey Smith G, Bowden J. Robust inference in summary data Mendelian randomization via the zero modal pleiotropy assumption. Int J Epidemiol. 2017;46:1985–1998.

13. Qi G, Chatterjee N. Mendelian randomization analysis using mixture models for robust and efficient estimation of causal effects. Nature Communications. 2019;10:1941.

14. Burgess S, Foley CN, Allara E, Staley JR, Howson JMM. A robust and efficient method for Mendelian randomization with hundreds of genetic variants: unravelling mechanisms linking HDL-cholesterol and coronary heart disease. bioRxiv. 2019 566851.

15. Bulik-Sullivan BK, Loh P-R, Finucane HK et al. LD Score regression distinguishes confounding from polygenicity in genome-wide association studies. Nat Genet. 2015;47:291–291.

16. Lee SH, Yang J, Goddard ME, Visscher PM, Wray NR. Estimation of pleiotropy between complex diseases using SNP-derived genomic relationships and restricted maximum likelihood. Bioinformatics. 2012;28:2540–2542.

17. Speed D, Hemani G, Johnson MR, Balding DJ. Improved heritability estimation from genome-wide SNPs. The American Journal of Human Genetics. 2012;91:1011–1021.

18. Speed D, Balding DJ. SumHer better estimates the SNP heritability of complex traits from summary statistics. Nat Genet. 2019;51:277.

19. Yang J, Benyamin B, McEvoy BP et al. Common SNPs explain a large proportion of the heritability for human height. Nat Genet. 2010;42:565.

20. Stephens M. False discovery rates: a new deal. Biostatistics. 2016;18:275–294.

21. Zeng J, De Vlaming R, Wu Y et al. Signatures of negative selection in the genetic architecture of human complex traits. Nat Genet. 2018;50:746.

22. Zhang Y, Qi G, Park J-H, Chatterjee N. Estimation of complex effect-size distributions using summary-level statistics from genome-wide association studies across 32 complex traits. Nat Genet. 2018;50:1318.

23. Zhu X, Stephens M. Large-scale genome-wide enrichment analyses identify new trait-associated genes and pathways across 31 human phenotypes. Nature communications. 2018;9:4361.

24. Burgess S, Butterworth A, Thompson SG. Mendelian randomization analysis with multiple genetic variants using summarized Data. Genet Epidemiol. 2013;37:658–665.

25. Burgess S, Bowden J, Dudbridge F, Thompson SG. Robust instrumental variable methods using multiple candidate instruments with application to Mendelian randomization. arXiv preprint 160603729. 2016

26. Burgess S, Zuber V, Gkatzionis A, Foley CN. Modal-based estimation via heterogeneity-penalized weighting: model averaging for consistent and efficient estimation in Mendelian randomization when a plurality of candidate instruments are valid. Int J Epidemiol. 2018 dyy080.

27. Loh P-R, Tucker G, Bulik-Sullivan BK et al. Efficient Bayesian mixed-model analysis increases association power in large cohorts. Nat Genet. 2015;47:284.

28. Slob EAW, Burgess S. A Comparison Of Robust Mendelian Randomization Methods Using Summary Data. BioRxiv. 2019 577940.

29. Zhao Q, Wang J, Bowden J, Small DS. Statistical inference in two-sample summary-data Mendelian randomization using robust adjusted profile score. arXiv preprint 180109652. 2018

30. Bowden J, Del Greco M F, Minelli C, Davey Smith G, Sheehan N, Thompson J. A framework for the investigation of pleiotropy in two-sample summary data Mendelian randomization. Stat Med. 2017;36:1783–1802.

31. Sinnott-Armstrong N, Tanigawa Y, Amar D et al. Genetics of 38 blood and urine biomarkers in the UK Biobank. BioRxiv. 2019 660506.

32. Mahajan A, Taliun D, Thurner M et al. Fine-mapping type 2 diabetes loci to single-variant resolution using high-density imputation and islet-specific epigenome maps. Nat Genet. 2018;50:1505.

