## Supplementary Figures and Tables for "A Comprehensive Evaluation of Methods for Mendelian Randomization Using Realistic Simulations and an Analysis of 38 Biomarkers for Risk of Type-2 Diabetes"

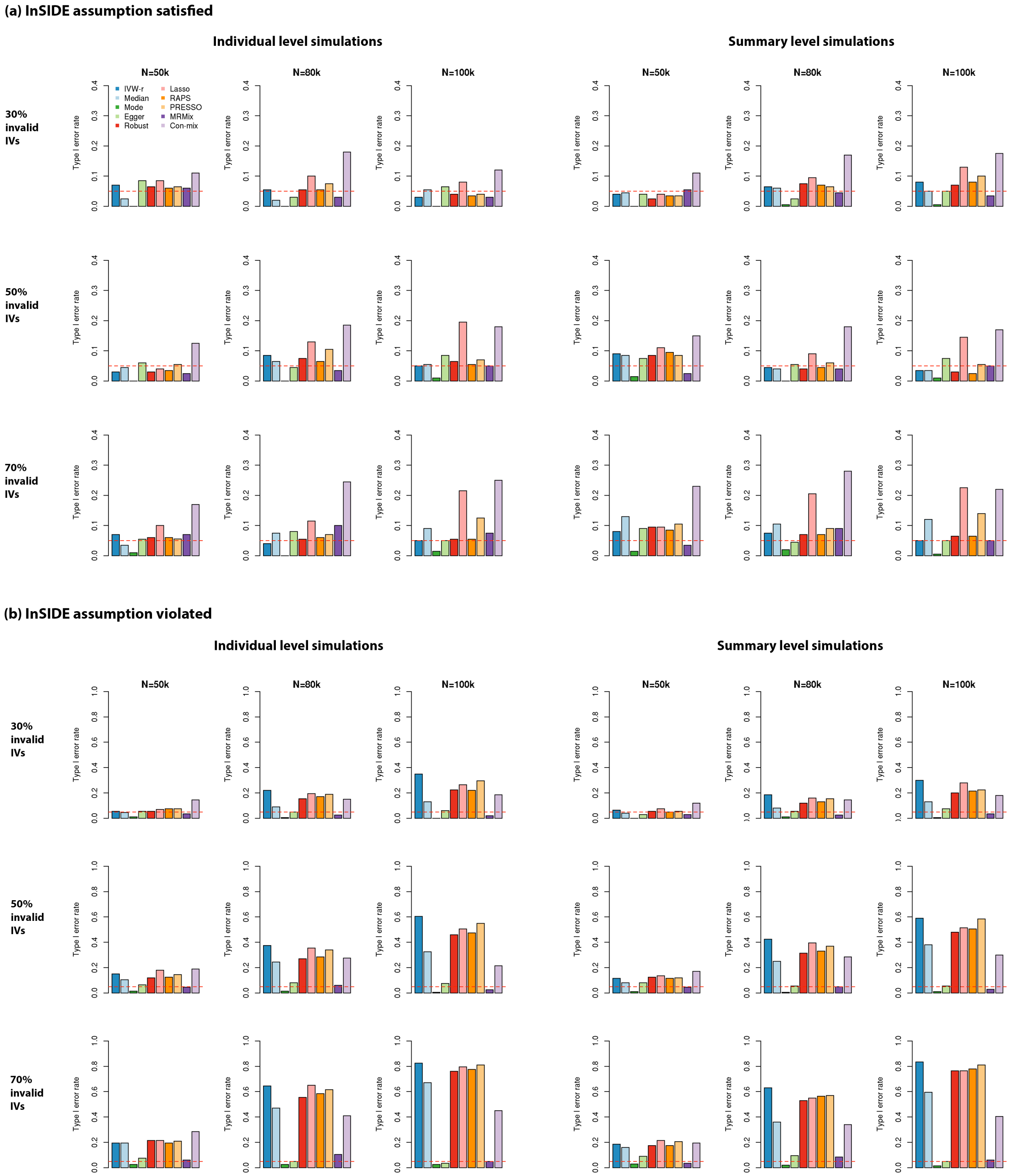


**Supplementary Figure 1. Comparison of type I error rates of different methods using individual- and summary-level data simulations.** Data are simulated under balance pleiotropy with or without InSIDE assumption satisfied. Sample size of the study associated with $X$ is $N$; sample size of the study associated with $Y$ is $N/2$. The red dashed line is the nominal significance threshold 0.05. Empirical type I error rates are calculated over 200 simulations.

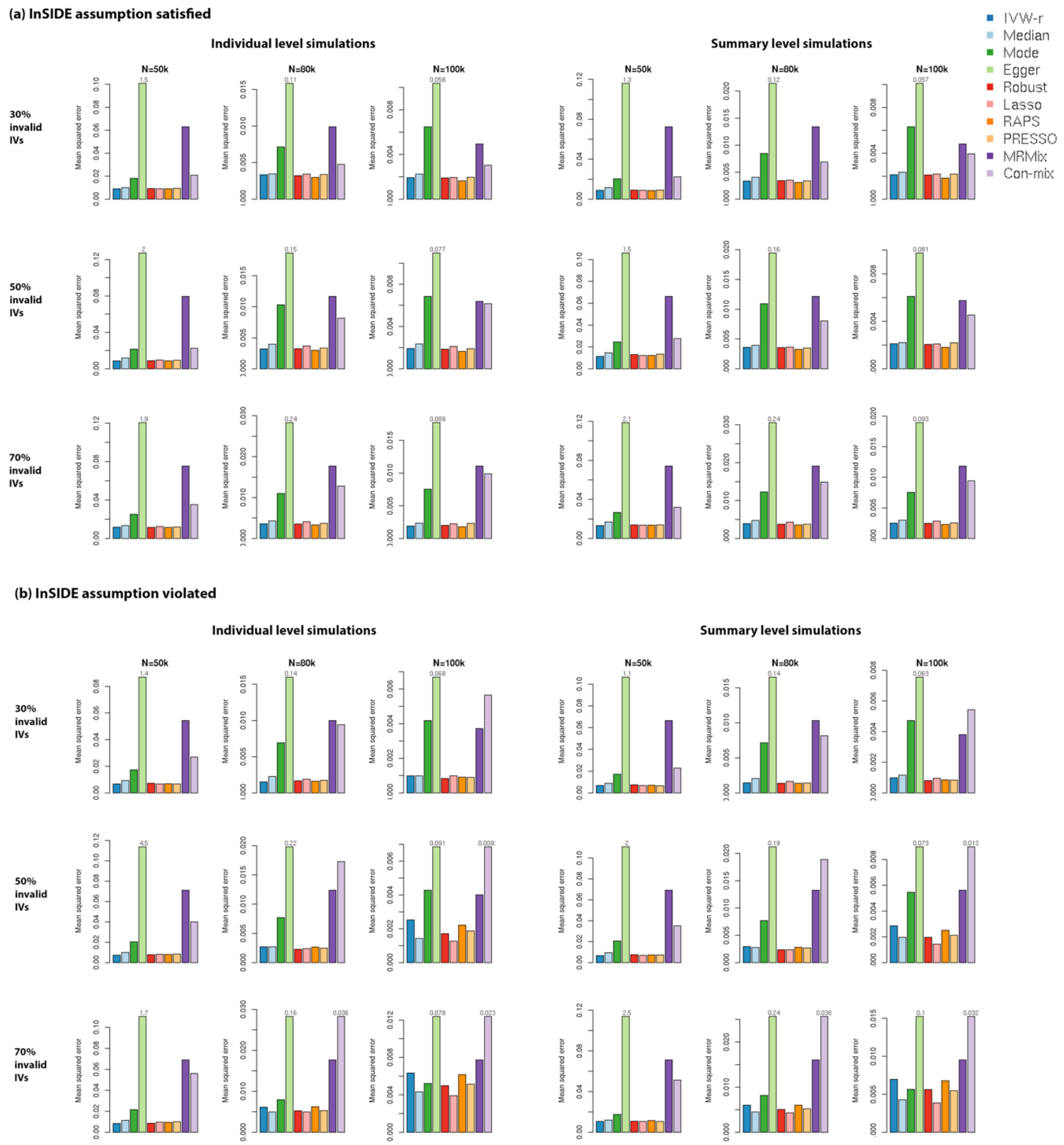


**Supplementary Figure 2. Comparison of mean squared errors (MSE) of different MR methods using individual- and summary-level simulations.** Data are simulated assuming a true causal effect of 0.2 and balanced pleiotropy, with or without InSIDE assumption. Sample size of the study associated with $X$ is $N$; sample size of the study associated with $Y$ is $N/2$. Bars higher than the upper limit of the panel are truncated and marked with the true value. All MSEs are computed over 200 simulations.

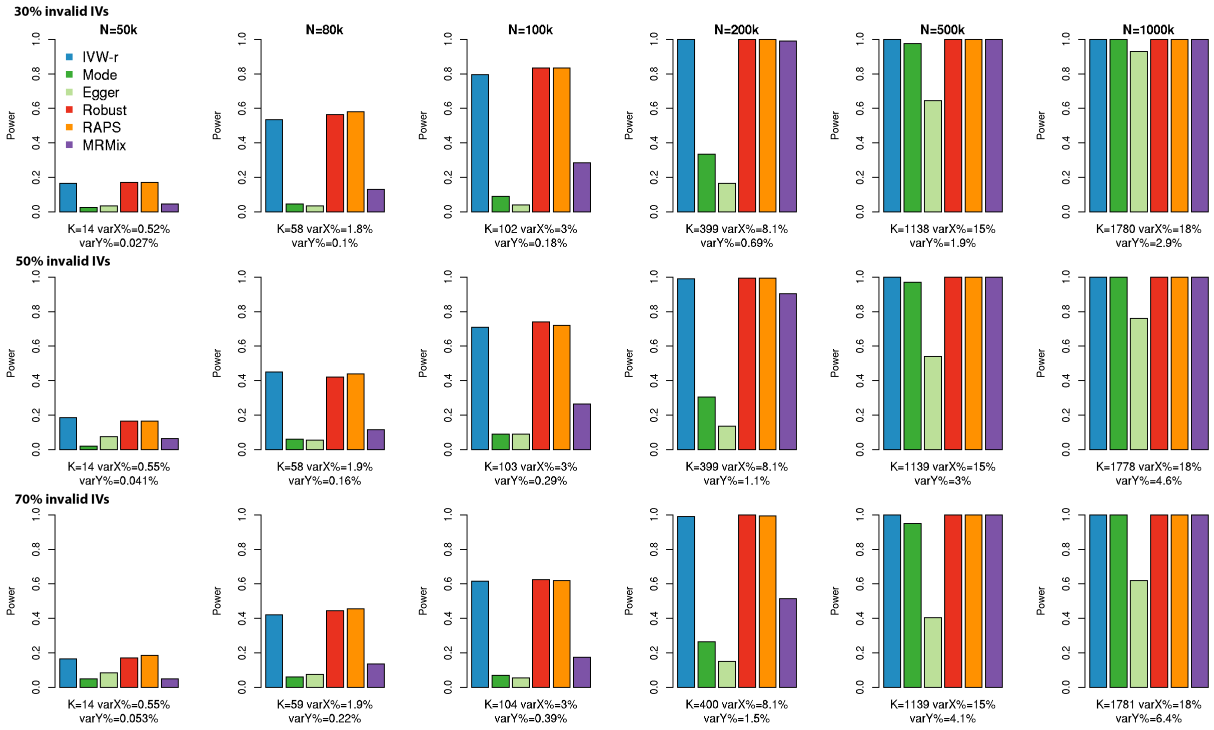


**Supplementary Figure 3**. **Power of the methods that have well controlled or moderately inflated type I error rates in simulations with balanced pleiotropy and InSIDE assumption satisfied.** Data are simulated assuming a true causal effect of 0.1. Power is reported over 200 simulations. Sample size of the study associated with $X$ is $N$; sample size of the study associated with $Y$ is $N/2$. $K$: the average number of IVs, defined as the SNPs which reach genome-wide significance (z-test $p < 5\times10^{-8}$) in the study associated with $X$; varX%: average percentage of variance of $X$ explained by IVs; varY%: average percentage of variance of $Y$ explained by IVs.

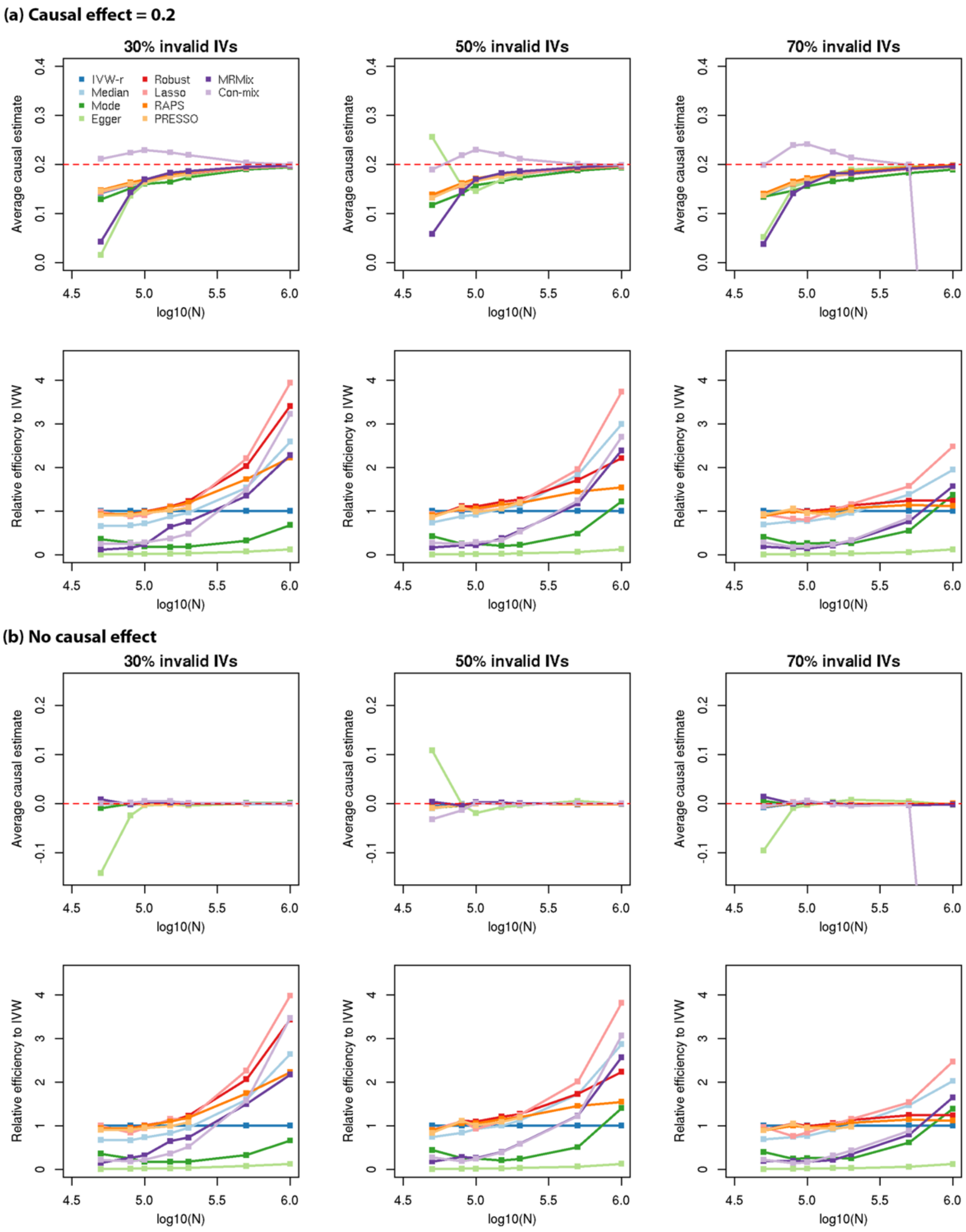


**Supplementary Figure 4. Mean estimates and standard error of method relative to that of IVW-r in simulations with balanced pleiotropy and InSIDE assumption satisfied.** Sample size of the study associated with $X$ is $N$; sample size of the study associated with $Y$ is $N/2$. The red dashed line is the true causal effect. All results are reported over 200 simulations.

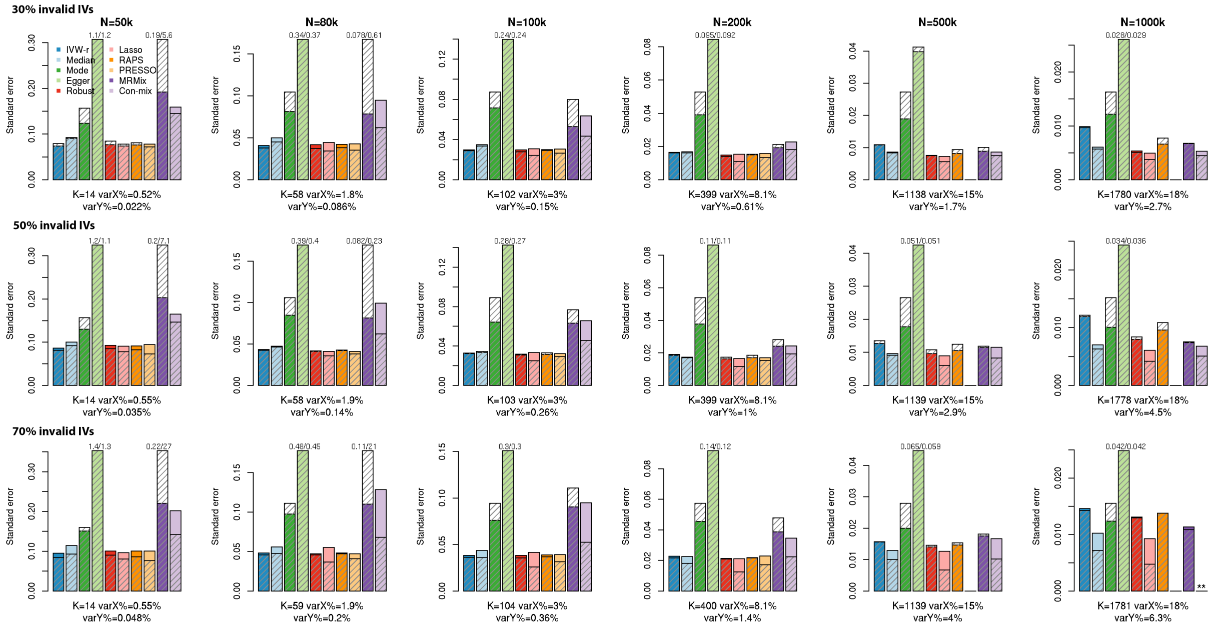


**Supplementary Figure 5. Empirical and estimated standard errors in simulations with balanced pleiotropy, InSIDE assumption satisfied and no causal effects.** The colored bars are the standard deviations of causal effect estimates over 200 simulations (empirical standard error); the shaded bars are the average of estimated standard errors (SE) over 200 simulations. The SE estimate is considered conservative if the colored bar is lower than the superimposed shaded bar; and anticonservative vice versa. Sample size of the study associated with $X$ is $N$; sample size of the study associated with $Y$ is $N/2$. $K$: the average number of IVs, defined as the SNPs which reach genome-wide significance (z-test $p < 5\times10^{-8}$) in the study associated with $X$; varX%: average percentage of variance of $X$ explained by IVs; varY%: average percentage of variance of $Y$ explained by IVs. Bars higher than the upper limit of the panel are truncated and marked with the true value (empirical SE/average estimated SE). When $N = 1000k$ and 70% of the IVs are invalid, Con-mix estimates the causal effect to be -1 in all simulations; hence the SE is not shown and marked with $**$.

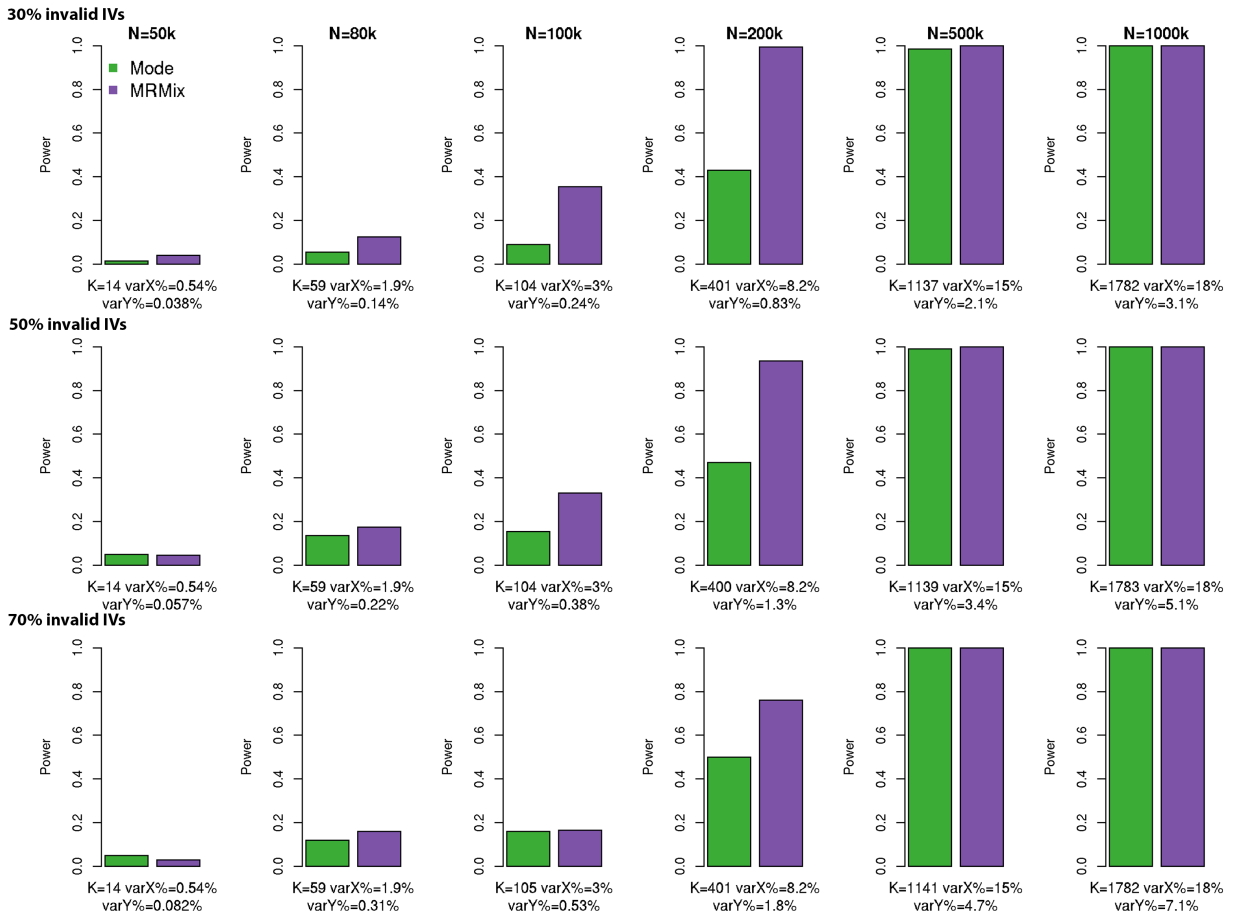


**Supplementary Figure 6**. **Power of weighted mode and MRMix in simulations with balanced pleiotropy and InSIDE assumption violated.** Data are simulated assuming a true causal effect of 0.1. Sample size of the study associated with $X$ is $N$; sample size of the study associated with $Y$ is $N/2$. $K$: the average number of IVs, defined as the SNPs which reach genome-wide significance (z-test $p < 5\times10^{-8}$) in the study associated with $X$; varX%: average percentage of variance of $X$ explained by IVs; varY%: average percentage of variance of $Y$ explained by IVs. All results are reported over 200 simulations.

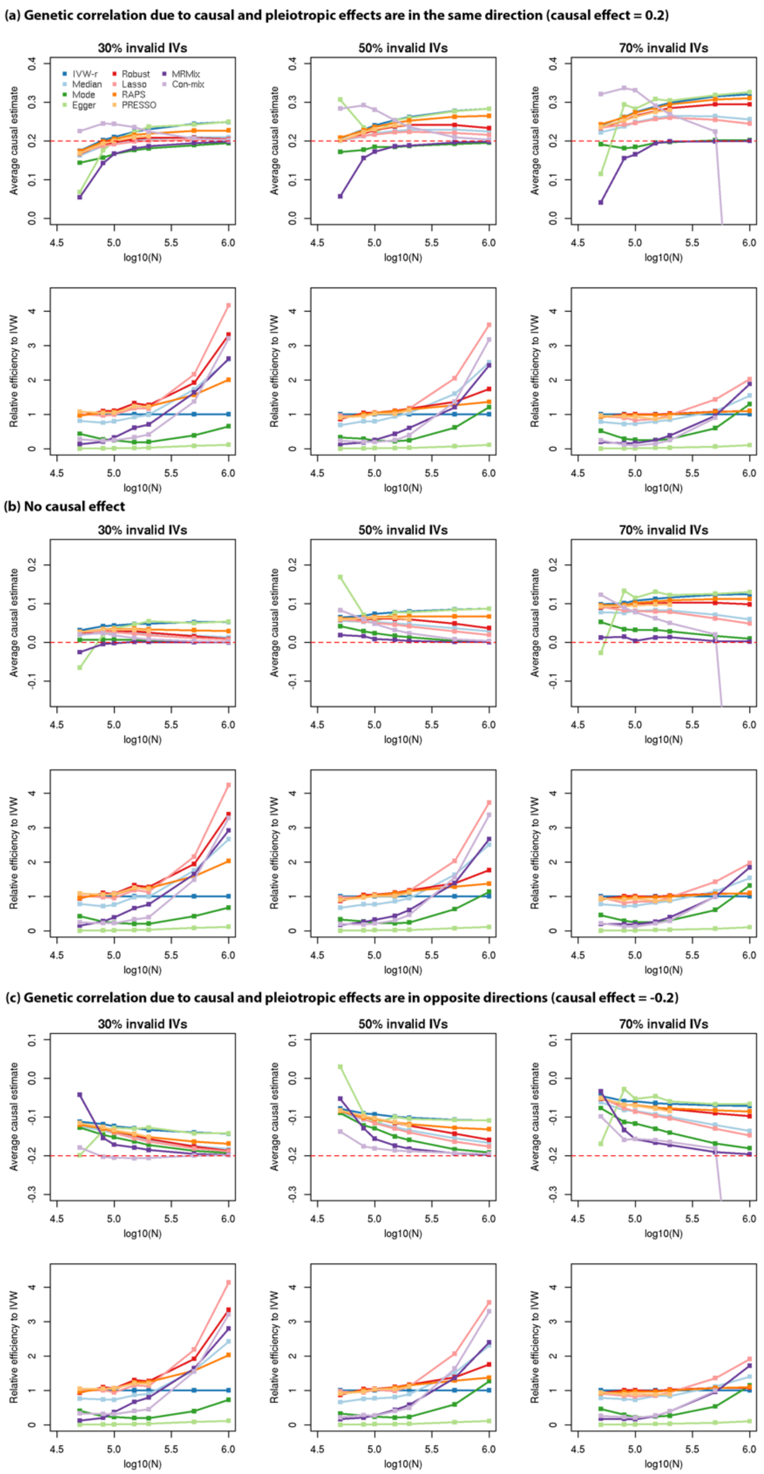


**Supplementary Figure 7. Mean estimate and standard errors relative to that of IVW-r in simulations with balanced pleiotropy and InSIDE assumption violated.** Sample size of the study associated with $X$ is $N$; sample size of the study associated with $Y$ is $N/2$. The red dashed line is the true causal effect. All results are reported over 200 simulations.

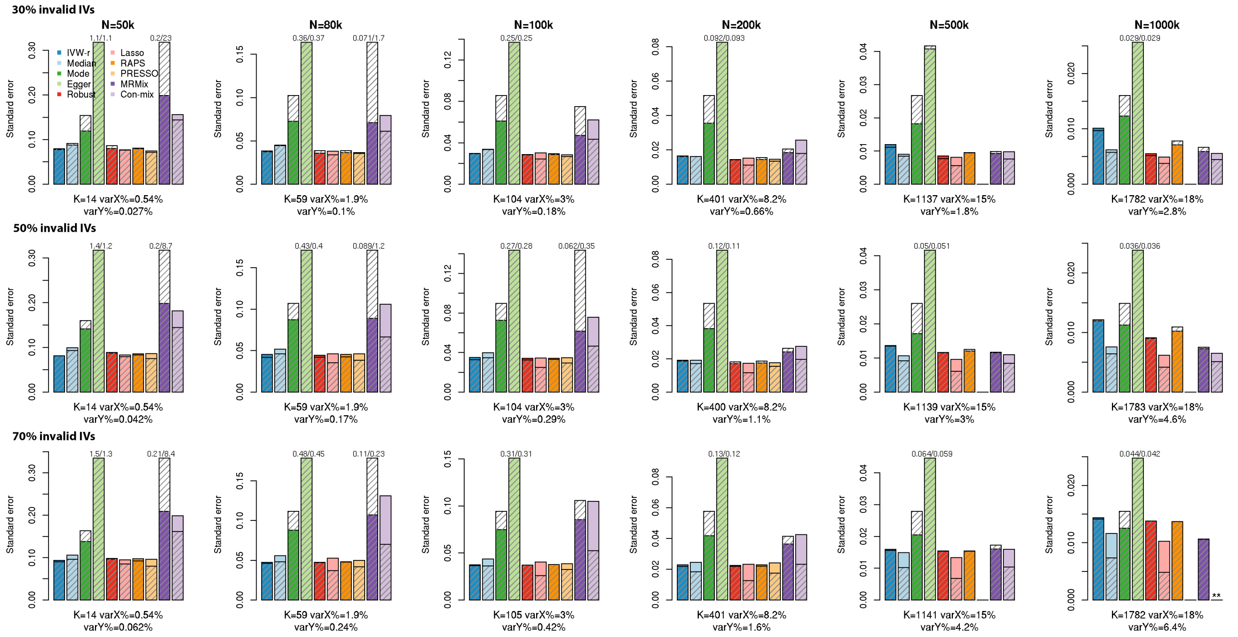


**Supplementary Figure 8. Empirical and estimated standard errors in simulations with balanced pleiotropy, InSIDE assumption violated and no causal effects.** The colored bars are the standard deviations of causal effect estimates over 200 simulations (empirical standard error); the shaded bars are the average of estimated standard errors (SE) over 200 simulations. The SE estimate is considered conservative if the colored bar is lower than the superimposed shaded bar; and anticonservative vice versa. Sample size of the study associated with $X$ is $N$; sample size of the study associated with $Y$ is $N/2$. $K$: the average number of IVs, defined as the SNPs which reach genome-wide significance (z-test $p < 5\times10^{-8}$) in the study associated with $X$; varX%: average percentage of variance of $X$ explained by IVs; varY%: average percentage of variance of $Y$ explained by IVs. Bars higher than the upper limit of the panel are truncated and marked with the true value (empirical SE/average estimated SE). When $N = 1000k$ and 70% of the IVs are invalid, Con-mix estimates the causal effect to be -1 in all simulations; hence the SE is not shown and marked with **.

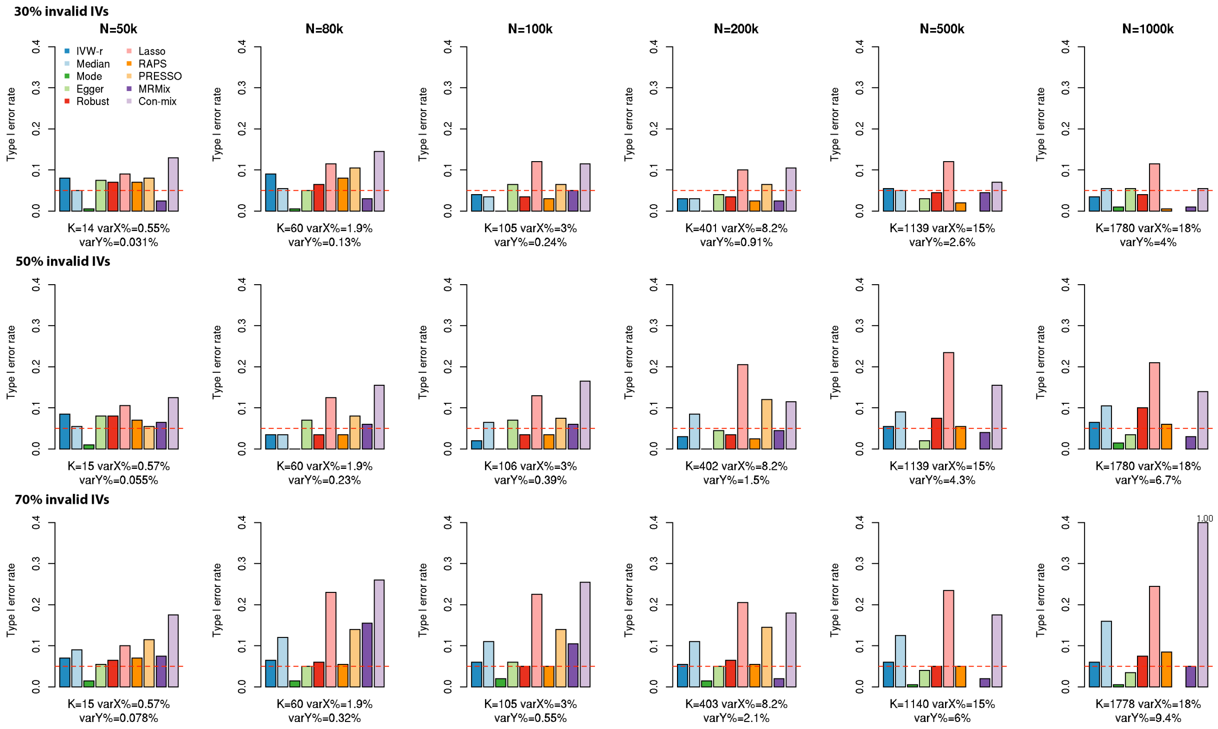


**Supplementary Figure 9. Type I error rate in simulations with directional pleiotropy and InSIDE assumption satisfied.** Sample size of the study associated with $X$ is $N$; sample size of the study associated with $Y$ is $N/2$. $K$: the average number of IVs, defined as the SNPs which reach genome-wide significance (z-test $p < 5\times10^{-8}$) in the study associated with $X$; varX%: average percentage of variance of $X$ explained by IVs; varY%: average percentage of variance of $Y$ explained by IVs. The red dashed line is the nominal significance threshold 0.05. Bars higher than the upper limit of the panel are truncated and marked with the true value. Empirical type I errors are reported over 200 simulations.

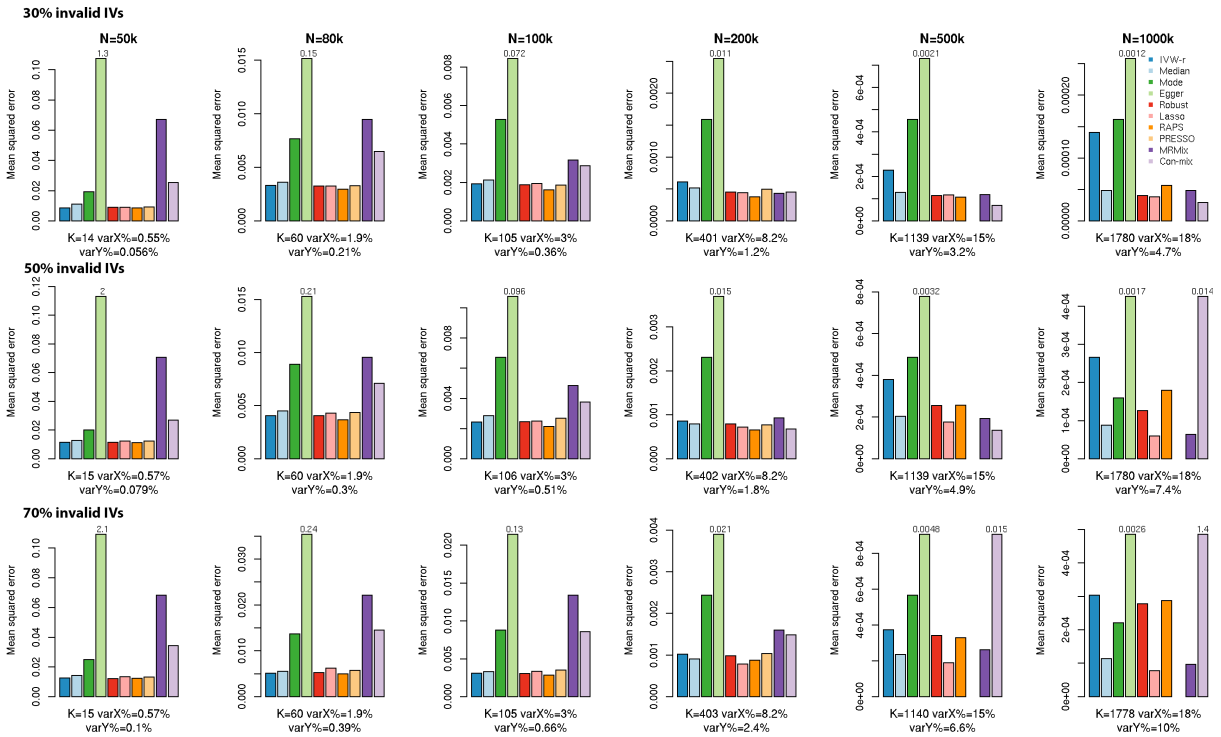


**Supplementary Figure 10. Mean squared error in simulations with directional pleiotropy and InSIDE assumption satisfied.** Data are simulated assuming a true causal effect of 0.2. Sample size of the study associated with $X$ is $N$; sample size of the study associated with $Y$ is $N/2$. $K$: the average number of IVs, defined as the SNPs which reach genome-wide significance (z-test $p < 5\times10^{-8}$) in the study associated with $X$; varX%: average percentage of variance of $X$ explained by IVs; varY%: average percentage of variance of $Y$ explained by IVs. Bars higher than the upper limit of the panel are truncated and marked with the true value. All results are reported over 200 simulations.

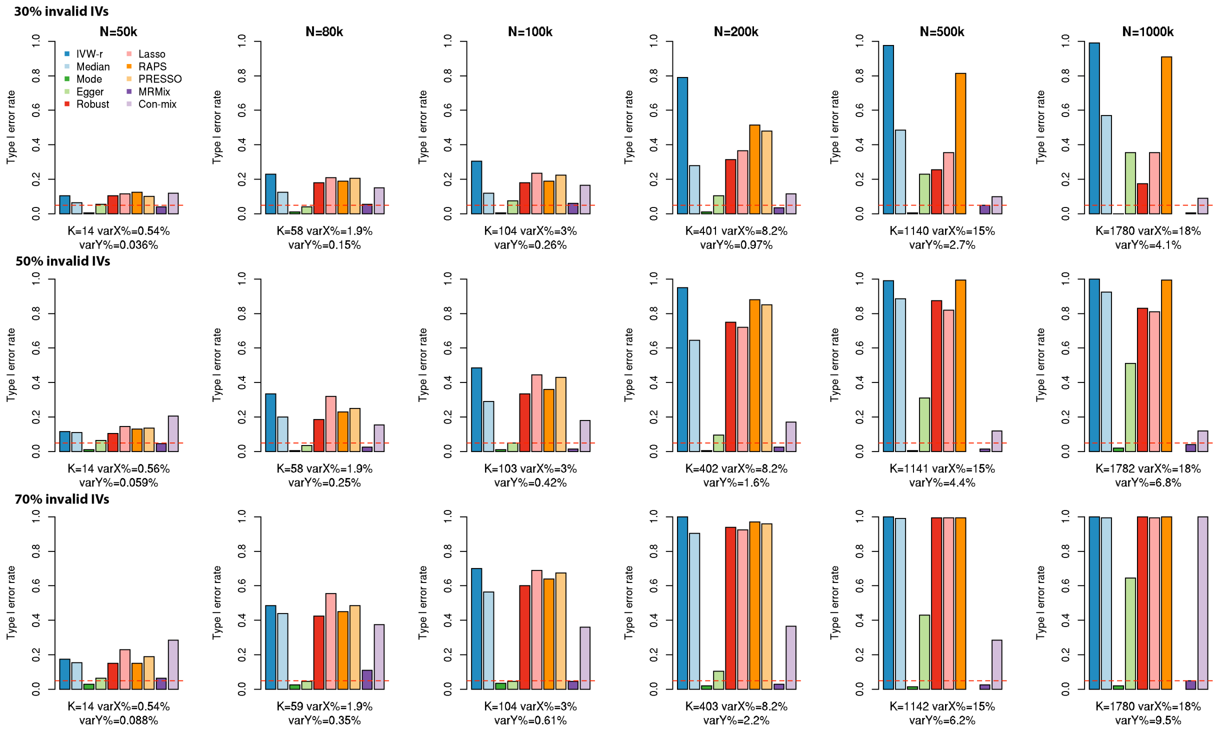


**Supplementary Figure 11. Type I error rate in simulations with directional pleiotropy and InSIDE assumption violated.** Sample size of the study associated with $X$ is $N$; sample size of the study associated with $Y$ is $N/2$. $K$: the average number of IVs, defined as the SNPs which reach genome-wide significance (z-test $p < 5\times10^{-8}$) in the study associated with $X$; varX%: average percentage of variance of $X$ explained by IVs; varY%: average percentage of variance of $Y$ explained by IVs. The red dashed line is the nominal significance threshold 0.05. Empirical type I errors are reported over 200 simulations.

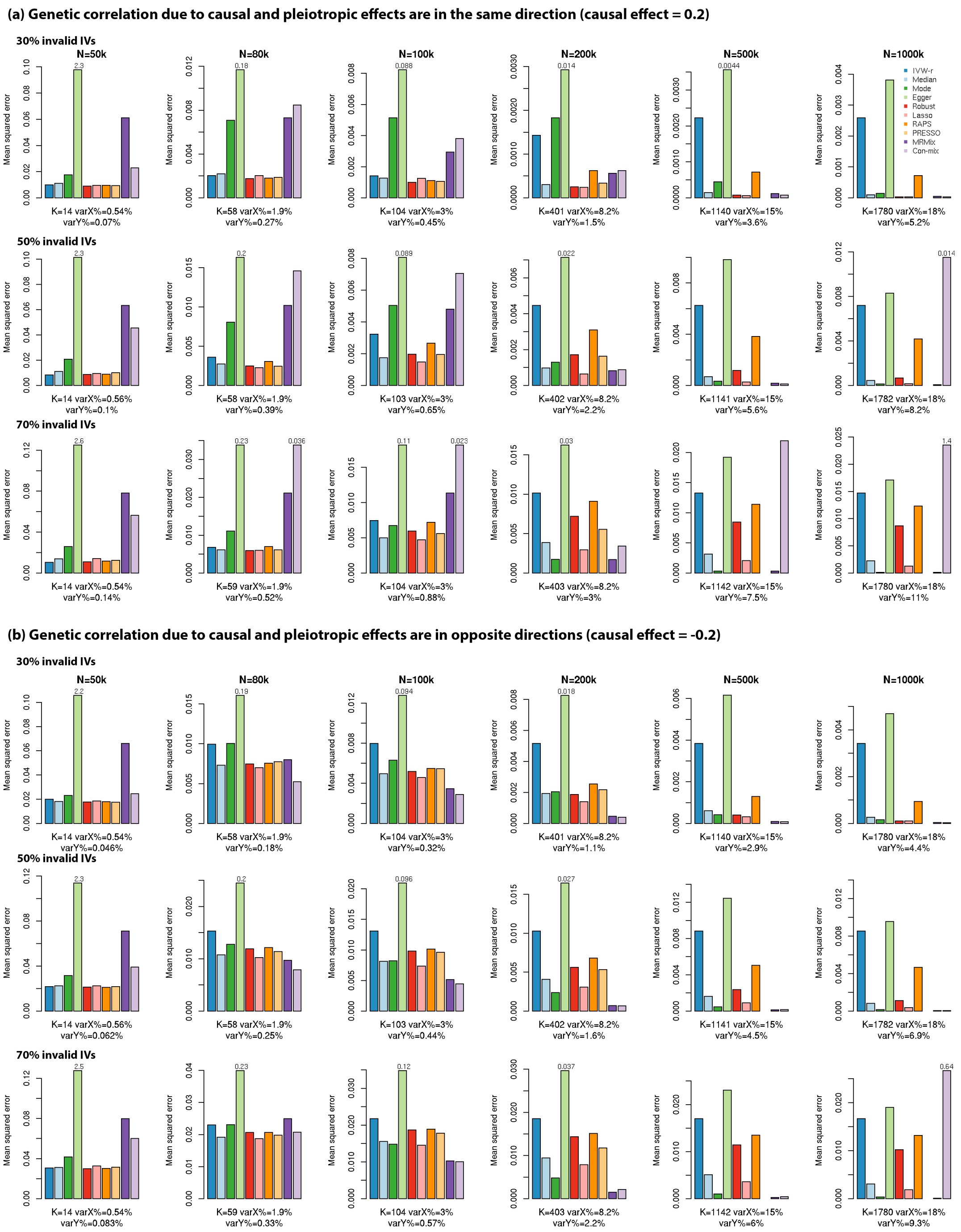


**Supplementary Figure 12. Mean squared error in simulations with directional pleiotropy and InSIDE assumption violated.** Sample size of the study associated with $X$ is $N$; sample size of the study associated with $Y$ is $N/2$. $K$: the average number of IVs, defined as the SNPs which reach genome-wide significance (z-test $p < 5\times10^{-8}$) in the study associated with $X$; varX%: average percentage of variance of $X$ explained by IVs; varY%: average percentage of variance of $Y$ explained by IVs. Bars higher than the upper limit of the panel are truncated and marked with the true value. All results are reported over 200 simulations.

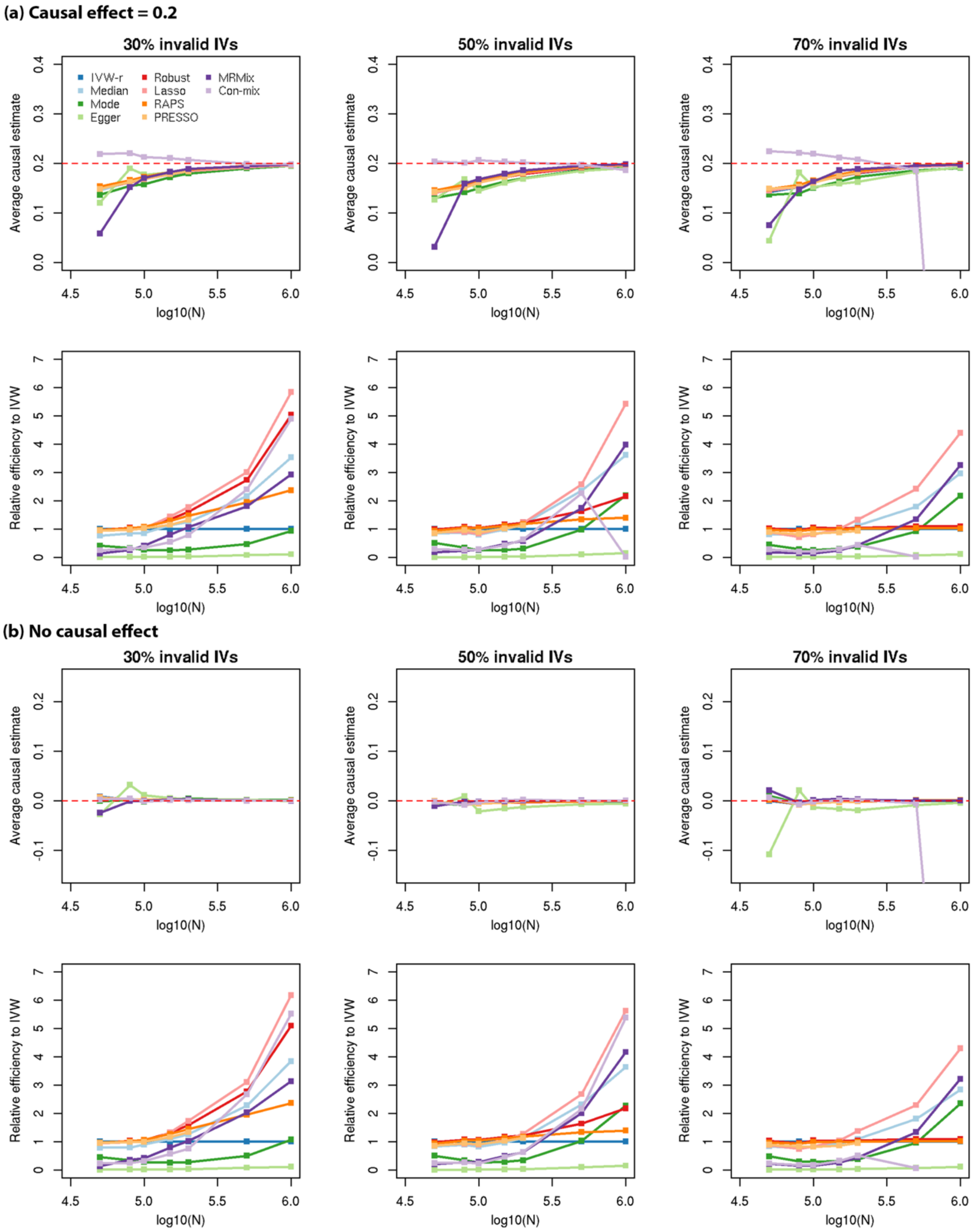


**Supplementary Figure 13. Mean estimate and standard errors relative to that of IVW-r in simulations with directional pleiotropy and InSIDE assumption satisfied.** Sample size of the study associated with $X$ is $N$; sample size of the study associated with $Y$ is $N/2$. The red dashed line is the true causal effect. All results are reported over 200 simulations.

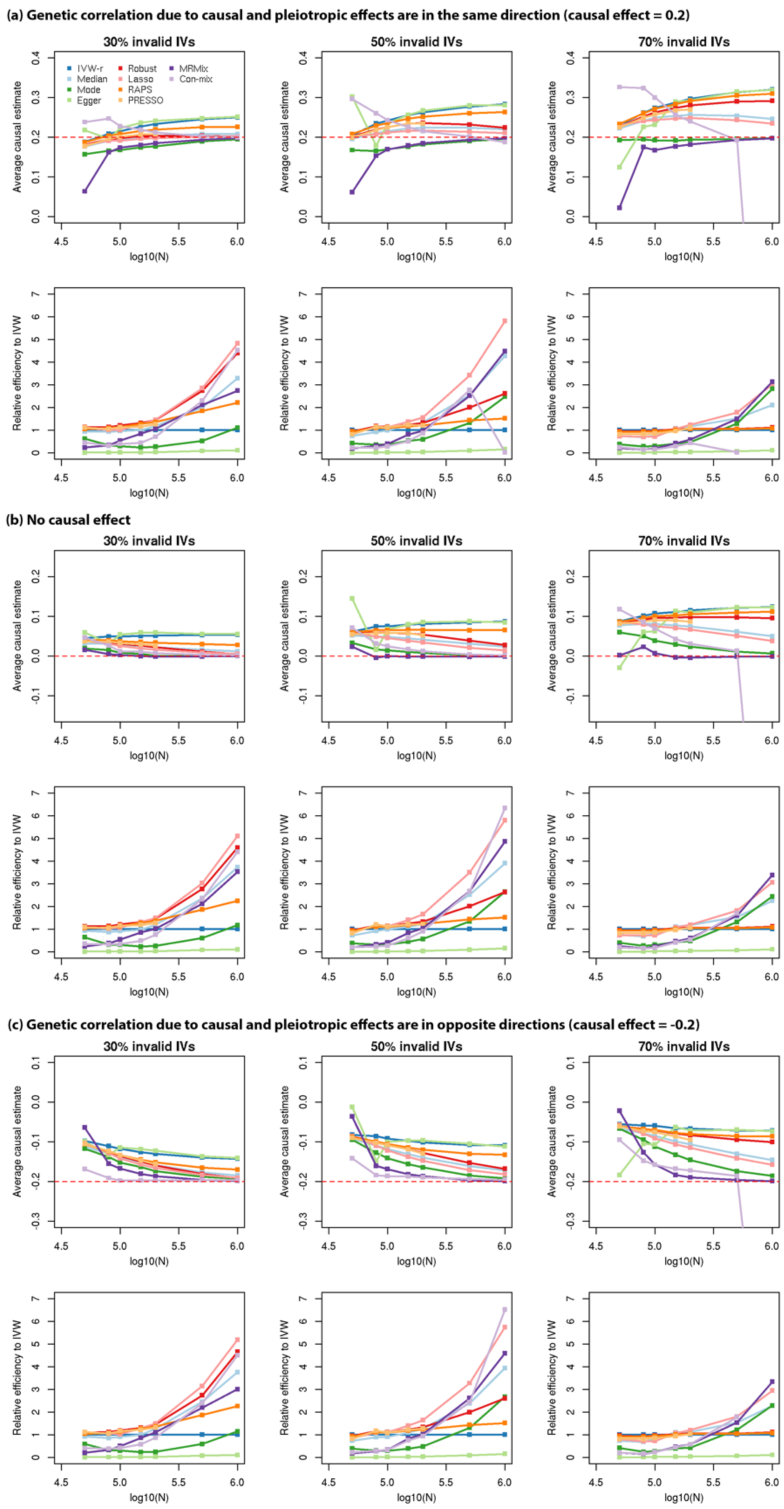


**Supplementary Figure 14. Mean estimates and standard errors relative to that of IVW-r in simulations with directional pleiotropy and InSIDE assumption violated.** Sample size of the study associated with $X$ is $N$; sample size of the study associated with $Y$ is $N/2$. The red dashed line is the true causal effect. All results are reported over 200 simulations.

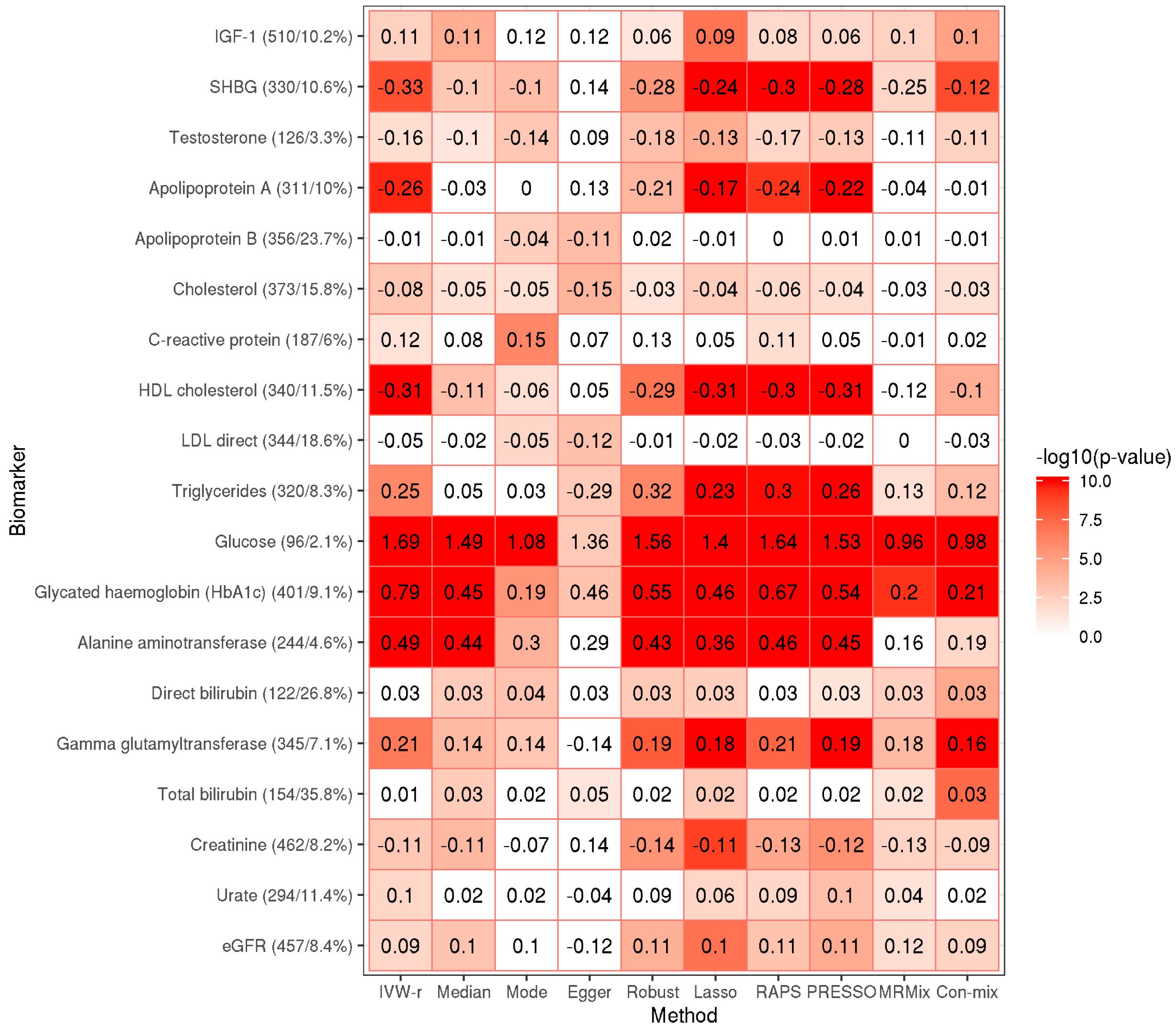


**Supplementary Figure 15. MR estimates and corresponding p-values of causal effects from UK Biobank blood and urine biomarkers to type 2 diabetes**. The numbers in each cell are the log-OR of type 2 diabetes (T2D) per SD increase in the exposure. We present the results for biomarkers that satisfy three criteria: 1) >25 IVs 2) IVs explain >1% variance of the exposure 3) have a causal effect on T2D at $p < 0.005$ by at least one method. The white cells correspond to p-values $< 0.05$, and those in color have p-values $\geq 0.05$; p-values that are $< 1\times10^{-10}$ are truncated at $1\times10^{-10}$. Each biomarker name is followed by (number of IVs/percentage of variance of biomarker explained by the IVs). Con-mix originally does not provide standard errors (SE) or p-values, but to compare with other methods, we first calculate an equivalent standard error, defined by (length of 95% confidence interval)/3.92, and then use the equivalent SE to calculate p-values.

### Supplementary Table 1. Simulation parameter settings

| **Setting** | **Causal effect** | **Effect of confounder** | **Mixture probabilities** | **Variance parameters** | **Directional pleiotropic effects** |
|---|---|---|---|---|---|
| Balanced pleiotropy, InSIDE satisfied [Model (M1), Methods] | 1) $\theta = 0.2$: genetic correlation due to causal and pleiotropic effects are in the same direction.<br><br>2) $\theta = 0$: no causal effects<br><br>3) $\theta = -0.2$: genetic correlation due to causal and pleiotropic effects are in opposite directions. | $\theta_{Ux} = \theta_{Uy} = 0.3$ | $\pi_3 = 0.01$<br>$\pi_4 = 0.97$<br><br>1) 30% invalid IVs<br>$\pi_1 = 0.14, \pi_2 = 0.06$<br>2) 50% invalid IVs<br>$\pi_1 = 0.1, \pi_2 = 0.1$<br>3) 70% invalid IVs<br>$\pi_1 = 0.06, \pi_2 = 0.14$ | $\sigma_x^2 = \sigma_y^2 = 5\times10^{-5}$ | NA |
| Balanced pleiotropy, InSIDE violated [Model (M2), Methods] | | | | $\sigma_x^2 = \sigma_y^2 = 5\times10^{-5}$<br>$\sigma_u^2 = 1\times10^{-4}$<br>$\tilde{\sigma}_x^2 = \tilde{\sigma}_y^2 = 4.1\times10^{-5}$ | NA |
| Directional pleiotropy, InSIDE satisfied [Model (M3), Methods] | | | | $\sigma_x^2 = \sigma_y^2 = 5\times10^{-5}$ | $\mu_y = 0.005.$ |
| Directional pleiotropy, InSIDE violated [Model (M4), Methods] | | | | $\sigma_x^2 = \sigma_y^2 = 5\times10^{-5}$<br>$\sigma_u^2 = 1\times10^{-4}$<br>$\tilde{\sigma}_x^2 = \tilde{\sigma}_y^2 = 4.1\times10^{-5}$ | |

**Supplementary Table 2. Variance of $X$ and $Y$ explained by direct effects of the instruments in Slob and Burgess simulation study under balanced pleiotropy and InSIDE assumption.** Here $\theta = 0.2$.

| Setting | Proportion of var(X) explained by direct effects of IVs (%) | Proportion of var(Y) explained by direct effects of IVs (%) |
|---|---|---|
| 10 IVs 30% invalid | 10 | 7.1 |
| 10 IVs 50% invalid | 10 | 11.2 |
| 10 IVs 70% invalid | 10 | 15.1 |
| 30 IVs 30% invalid | 10 | 18.6 |
| 30 IVs 50% invalid | 10 | 27.5 |
| 30 IVs 70% invalid | 10 | 34.7 |
| 100 IVs 30% invalid | 10 | 43.2 |
| 100 IVs 50% invalid | 10 | 55.9 |
| 100 IVs 70% invalid | 10 | 63.9 |
| 500 IVs 30% invalid | 10 | 79.2 |
| 500 IVs 50% invalid | 10 | 86.4 |
| 500 IVs 70% invalid | 10 | 89.9 |

Reference: Slob, E. A. W. & Burgess, S. A Comparison of Robust Mendelian Randomization Methods Using Summary Data. *BioRxiv* 577940 (2019)

**Supplementary Table 3. Ten MR methods studied in this paper and corresponding software.**

| Method | R package | Version | Function and tuning parameter | Reference |
|---|---|---|---|---|
| IVW-r | MendelianRandomization | 0.4.0 | mr_ivw(): default settings | [1] |
| Weighted median | MendelianRandomization | 0.4.0 | mr_median(): default settings | [2] |
| Weighted mode | MendelianRandomization | 0.4.0 | mr_mbe(): default settings | [3] |
| Egger regression | MendelianRandomization | 0.4.0 | mr_egger(): default settings | [4] |
| MR-Robust | MendelianRandomization | 0.4.0 | mr_ivw(mr.obj, "random", robust = TRUE) | [5] |
| MR-Lasso | R code provided by Slob & Burgess [6] | NA | MR_lasso(): default settings | [5] |
| MR-RAPS | mr.raps | NA | mr.raps.overdispersed.robust(b_exp, b_out, se_exp, se_out, loss.function = "huber", k = 1.345, initialization = c("l2"), suppress.warning = FALSE, diagnostics = FALSE, niter = 20, tol = .Machine$double.eps^0.5) | [7] |
| MR-PRESSO | MRPRESSO | NA | mr_presso(BetaOutcome = "by", BetaExposure = "bx", SdOutcome = "byse", SdExposure = "bxse", OUTLIERtest = TRUE, DISTORTIONtest = TRUE, data = presso.df, NbDistribution = 3000, SignifThreshold = 0.05) | [8] |
| MRMix | MRMix | NA | MRMix(): theta_temp_vec = seq(-0.5,0.5,by=0.01), other settings are default. | [9] |
| Contamination mixture | MendelianRandomization | 0.4.0 | mr_conmix(): default settings | [10] |

**References for Supplementary Tables**


1. Bowden, J. et al. A framework for the investigation of pleiotropy in two-sample summary data Mendelian randomization. *Stat Med* **36**, 1783-1802 (2017).
2. Bowden, J., Davey, S. G., Haycock, P. C. & Burgess, S. Consistent estimation in Mendelian randomization with some invalid instruments using a weighted median estimator. *Genet. Epidemiol.* **40**, 304-314 (2016).
3. Hartwig, F. P., Davey Smith, G. & Bowden, J. Robust inference in summary data Mendelian randomization via the zero modal pleiotropy assumption. *Int J Epidemiol* **46**, 1985-1998 (2017).
4. Bowden, J., Davey Smith, G. & Burgess, S. Mendelian randomization with invalid instruments: effect estimation and bias detection through Egger regression. *Int J Epidemiol* **44**, 512-525 (2015).
5. Burgess, S., Bowden, J., Dudbridge, F. & Thompson, S. G. Robust instrumental variable methods using multiple candidate instruments with application to Mendelian randomization. *arXiv preprint arXiv:1606.03729* (2016).
6. Slob, E. A. W. & Burgess, S. A Comparison Of Robust Mendelian Randomization Methods Using Summary Data. *BioRxiv* 577940 (2019).
7. Zhao, Q., Wang, J., Bowden, J. & Small, D. S. Statistical inference in two-sample summary-data Mendelian randomization using robust adjusted profile score. *arXiv preprint arXiv:1801.09652* (2018).
8. Verbanck, M., Chen, C.-Y., Neale, B. & Do, R. Detection of widespread horizontal pleiotropy in causal relationships inferred from Mendelian randomization between complex traits and diseases. *Nat Genet* **50**, 693 (2018).
9. Qi, G. & Chatterjee, N. Mendelian randomization analysis using mixture models for robust and efficient estimation of causal effects. *Nature Communications* **10**, 1941 (2019).
10. Burgess, S., Foley, C. N., Allara, E., Staley, J. R. & Howson, J. M. M. A robust and efficient method for Mendelian randomization with hundreds of genetic variants: unravelling mechanisms linking HDL-cholesterol and coronary heart disease. *bioRxiv* 566851 (2019).
