## Supplementary Notes for "A Comprehensive Evaluation of Methods for Mendelian Randomization Using Realistic Simulations and an Analysis of 38 Biomarkers for Risk of Type-2 Diabetes"

### 1. Details of Simulation Studies

We begin by introducing a few notations. Let $X$ denote the exposure, $Y$ denote the outcome and $U$ denote a potential confounder. Let $G_j$ denote the genotype of SNP $j$. We simulate data using the following model

$$U = \sum_{j=1}^{M} \phi_j G_j + \epsilon^U \quad (1)$$

$$X = \sum_{j=1}^{M} \gamma_j G_j + \theta_{Ux} U + \epsilon^X \quad (2)$$

$$Y = \sum_{j=1}^{M} \alpha_j G_j + \theta X + \theta_{Uy} U + \epsilon^Y \quad (3)$$

Here $\phi_j, \gamma_j, \alpha_j$ denote the direct effect of SNP $j$ on $U$, $X$ and $Y$, respectively. But unlike most previous studies that simulate only the selected instruments, we generate data for all common variants in the genome. The causal effect ($\theta$) of $X$ on $Y$ is the main parameter of interest. The effects of the confounder $U$ on $X$ and $Y$ are denoted by $\theta_{Ux}$ and $\theta_{Uy}$, respectively. The error terms $\epsilon^U$, $\epsilon^X$ and $\epsilon^Y$ are independent and normally distributed with mean 0. For convenience, we chose the variance of the error terms so that $U$, $X$ and $Y$ all have unit variance. We generate genotypes $G_j$ by first simulating $\tilde{G}_j$s independently from Binomial(2, 0.3) and then standardizing by $G_j = \frac{\tilde{G}_j - 2\times 0.3}{\sqrt{2\times 0.3\times(1-0.3)}}$ to make them have mean 0 and variance 1. They represent SNPs with minor allele frequency 0.3 after standardization. We generated data from model (1)-(3) using 200,000 independent SNPs as representative of all underlying common variants. We generate $\phi_j, \gamma_j, \alpha_j$ from mixture normal distributions as described in **Methods**, which have been shown to be appropriate for modeling effect-size distribution for complex traits in GWAS [1–4]. Under the above model, when the confounder $U$ has heritable component, the InSIDE assumption [5] is violated as direct and indirect effect of some SNPs on the outcome are correlated due to mediation by common factor $U$ .

*Simulating Data on Genome-wide Association Studies*

We simulate individual level data for independent genome-wide association studies for $X$ and $Y$ following the above model when sample size is not too large ($N \leq 100k$). We first conduct association analysis for each of the 200k SNPs with $X$ using data from the underlying GWAS. SNPs which reach genome-wide significance (p-value $< 5\times10^{-8}$) are then selected as instruments and then we analyze association of each of these SNPs with $Y$ using the underlying GWAS. However, for very large sample size, generation and analysis of individual level data can become computationally prohibitive and we simulate summary-level association statistics directly (addressed as *summary-level simulations*). We observe that the total effects of SNPs on $X$ and $Y$ are implied by model (1)-(3):

$$\beta_{jx} = \gamma_j + \theta_{Ux}\phi_j \quad (4)$$

$$\beta_{jy} = \alpha_j + \theta\beta_{jx} + \theta_{Uy}\phi_j \quad (5)$$

Thus, we simulate $\gamma_j, \phi_j$ and $\alpha_j$s as before and then directly generate summary statistics as $\hat{\beta}_{jx} = \beta_{jx} + N\left(0, \frac{1}{n_x}\right), \hat{\beta}_{jy} = \beta_{jy} + N\left(0, \frac{1}{n_y}\right)$ where the estimation error terms are generated with mean zero normal distribution with variances inversely proportional to $n_x$ and $n_y$, the sample size of the study associated with $X$ and $Y$, respectively. We explore sample sizes $(N)$ varying from 50k to 1000k with $n_x = N$ and $n_y = N/2$. Both individual and summary-level simulations are repeated 200 times.

*Summary of Simulation Results*

We calculate the type I error rate of all 10 methods at nominal significance threshold $p < 0.05$, as well as the power of the methods that have well controlled or only moderately inflated type I error. We also calculate the mean squared error (MSE) as

$$MSE = \frac{1}{200}\sum_{r=1}^{200}\left(\hat{\theta}_r - \theta^*\right)^2,$$

where $\theta^*$ is the true value of the causal effect. The MSE measures the accuracy of the point estimate combining bias and variance. We also use the mean and standard deviation of causal estimates across simulations to compare bias and efficiency separately. To investigate bias of the underlying standard error (SE) estimates in some of the MR methods, we compare the empirical standard deviation of the causal effect estimates and the average estimated SE calculated across simulations. For the contamination mixture method [6], we calculate the "standard error" as length of the 95% confidence set divided by $2\times1.96$, since the method generates confidence sets based on the likelihood ratio test and does not report a standard error.

**2. Unrealistic choice of parameters from an earlier simulation study**

We found previous studies for evaluation of MR methods have often not followed realistic model for genetic architecture of complex traits. In particular, a very recent study that evaluated a large number of methods for MR analysis following the basic setup described in (1)-(3), used highly unrealistic parameter settings [7]. The study, for example, assumes 10% of the variance of $X$ can be explained by the chosen instruments regardless of the number of instruments being 10, 30, 100 or 500. Results from existing GWAS reveal that complex traits tend to be extremely polygenic and only a very large number of variants can explain 10% of the variance of a trait. For example, latest GWAS has shown that more than 1000 SNPs are needed to explain 10% variance of a trait like BMI [8].

Further, the same study generated the direct effects of the chosen instruments on the outcome $(\alpha_j)$ to be so large that the instruments could explain more variability of $Y$ than that of $X$. For example, under the scenario with balanced pleiotropy they considered, the proportion of variance explained by direct effects of the instruments on $Y$ ranges from 20-35% for 30 SNPs and can go up to 90% for 500 IVs (**Supplementary Table 2**). Finally, the study fixes the sample size at 10,000 for both the exposure and the outcome – much smaller than the size of recent genome-wide association studies that have led to dozens or even hundreds of IVs for various traits. The study varies the number of instruments independently of the fixed sample size. In reality, sample size directly determines the number of instruments available and precision of their effects.

**3. Summary of Methods for Robust Mendelian Randomization Analysis**

We compare all nine methods investigated in the recent study indicated above [7]. In addition, we include the inverse-variance weighted method with multiplicative random effects (IVW-r) in comparison [9]. The methods can be classified into the following categories:

A. Location parameter of ratio estimates

- IVW-r: The IVW estimator with multiplicative random effects is a simple extension of the standard IVW. IVW-r computes the estimate of causal effect using the same weights as fixed-effect IVW, but incorporates an over-dispersion parameter into the variance to account for pleiotropy [9].

- Weighted median: The weighted median method takes the median of the ratio estimates after assigning to them probabilistic weights that are inversely proportional to their variances. The underlying assumption of that method is that >50% of the weight comes from valid IVs [10].
- Weighted mode: The weighted mode estimator takes the mode of the smoothed empirical density function of the ratio estimates, using the same weights as the weighted median approach. The method requires the ZEro Modal Pleiotropy Assumption (ZEMPA) [11].
- MR-Egger: Egger regression fits the regression model $\hat{\beta}_y = \theta\hat{\beta}_x + \theta_0$, where $\hat{\theta}$ is the estimated causal effect and $\hat{\theta}_0$ is the estimated directional pleiotropy. Since IVW estimator is in effect the slope of a regression model through the origin, Egger regression is a direct extension of the method by allowing an intercept term accounting for directional pleiotropy. The method requires the Instrument Strength Independent of Direct Effect (InSIDE) assumption [5].

B. Robust regression

- MR-Robust: IVW is performed by fitting the regression using MM-estimation, which consists of an initial S-estimate followed by an M-estimate of regression[12], combined with Tukey's bi-weight loss function [13].
- MR-Lasso: The IVW regression is augmented by adding SNP-specific intercept terms, which represents SNP-specific pleiotropy effects, and penalizing the intercept terms with L1 loss function [13].

- MR-RAPS: Profile likelihood can be used for MR when there is no horizontal pleiotropy. The MR Adjusted Profile Score method incorporates random effect and robust loss functions into the profile score to account for systematic and idiosyncratic pleiotropy [14].

C. Outlier detection and removal

- MR-PRESSO: The MR Pleiotropy Residual Sum and Outlier (MR-PRESSO) method uses the leave-one-out sum of squared residuals to detect global horizontal pleiotropy. It also detects and removes outliers that are in horizontal pleiotropy and conducts a distortion test of the influence of the invalid IVs. The MR-PRESSO outlier test requires that at least 50% of the variants are valid instruments and relies on the InSIDE assumption [15]. Due to long computational time, we only implements MR-PRESSO up to sample size $N = 200k$.

D. Mixture model approach

- MRMix: MRMix uses a mixture model for effect-size distribution assuming existence of a fraction of the genetic markers that are valid instruments. Causal effects are estimated based on a novel spike-detection algorithm: it fits the mixture model $\hat{\beta}_y - \theta\hat{\beta}_x \sim \pi_0 N(0, \sigma_0^2) + (1 - \pi_0)N(0, \sigma^2)$ and searches for the $\theta$ that maximizes the probability concentration at the null component $N(0, \sigma_0^2)$ corresponding to valid IVs. This approach requires ZEMPA and tends to be robust and efficient under large sample size [16].
- Contamination mixture (addressed as Con-mix for simplicity): The contamination mixture approach also uses a mixture model to characterize a cluster of valid IVs and a cluster of invalid IVs. Unlike MRMix which models the genetic effect size, Con-mix models the ratio

estimates using normal mixture with a pre-specified variance of the pleiotropic effects [6].

See **Supplementary Table 3** for the software and tuning parameters used to implement the methods above.

#### 4. MR Analysis for Biomarker effects on Type 2 diabetes

We applied the variety of available methods for MR analysis to investigate causal effect of 38 blood and urine biomarkers measures in the UK Biobank study [17] on the risk of type-2 diabetes. We accessed the summary statistics from recent analysis of the biomarkers in the UK Biobank study with the sample [17]. The biomarkers were adjusted for covariates including age, sex, genetic principal components, etc., and inverse normal transformed. The study analyzed unrelated UK Biobank individuals of white British ancestry that satisfy the following criteria: 1) not marked as outliers for heterozygosity and missing rates, 2) do not show putative sex chromosome aneuploidy 3) have at most 10 putative third degree relatives, which results in $N = 318\,984$ subjects. For each biomarker, we selected SNPs that are associated with each biomarker at $p < 5\times10^{-8}$ and further removed SNPs with $MAF \leq 0.01$ or in the MHC region. We restricted the analysis to those biomarkers which have at least 25 associated instruments and the Instruments explain at least 1% of the biomarker variance.

On the outcome side, we accessed summary statistics from the largest GWAS on type 2 diabetes [18]. The study consists of 74 124 T2D cases and 824 006 controls. Summary statistics for the biomarkers were merged with the summary statistics for the outcome, palindromic SNPs were removed, then LD clumping was performed at $r^2 < 0.1$ to remove the correlated SNPs. Lastly, for both the exposure and the outcome the estimates of genetic effects ($\hat{\beta}$) and the standard errors (SEs) were multiplied by $\sqrt{2 \times MAF(1 - MAF)}$ to transform them into the standardized scale. After the standardization the SEs for the genetic effects on biomarkers are roughly equal to $\frac{1}{\sqrt{N}}$, where $N$ is the sample size of the UK Biobank GWAS on biomarkers. This indicates that the biomarker measurements have been standardized to have unit variance, hence the MR estimates can be interpreted as log-OR of T2D per standard deviation (SD) unit increase in the exposure.

**References for Supplementary Notes**


1. Stephens M. False discovery rates: a new deal. Biostatistics. 2016;18:275-294.
2. Zeng J, De Vlaming R, Wu Y *et al.* Signatures of negative selection in the genetic architecture of human complex traits. Nat Genet. 2018;50:746.
3. Zhang Y, Qi G, Park J-H, Chatterjee N. Estimation of complex effect-size distributions using summary-level statistics from genome-wide association studies across 32 complex traits. Nat Genet. 2018;50:1318.
4. Zhu X, Stephens M. Large-scale genome-wide enrichment analyses identify new trait-associated genes and pathways across 31 human phenotypes. Nature communications. 2018;9:4361.
5. Bowden J, Davey Smith G, Burgess S. Mendelian randomization with invalid instruments: effect estimation and bias detection through Egger regression. Int J Epidemiol. 2015;44:512-525.
6. Burgess S, Foley CN, Allara E, Staley JR, Howson JMM. A robust and efficient method for Mendelian randomization with hundreds of genetic variants: unravelling mechanisms linking HDL-cholesterol and coronary heart disease. bioRxiv. 2019566851.
7. Slob EAW, Burgess S. A Comparison Of Robust Mendelian Randomization Methods

Using Summary Data. BioRxiv. 2019577940.

8. Yengo L, Sidorenko J, Kemper KE *et al.* Meta-analysis of genome-wide association studies for height and body mass index in~ 700000 individuals of European ancestry. Hum Mol Genet. 2018;27:3641-3649.

9. Bowden J, Del Greco M F, Minelli C, Davey Smith G, Sheehan N, Thompson J. A framework for the investigation of pleiotropy in two-sample summary data Mendelian randomization. Stat Med. 2017;36:1783-1802.

10. Bowden J, Davey SG, Haycock PC, Burgess S. Consistent estimation in Mendelian randomization with some invalid instruments using a weighted median estimator. Genet Epidemiol. 2016;40:304-314.

11. Hartwig FP, Davey Smith G, Bowden J. Robust inference in summary data Mendelian randomization via the zero modal pleiotropy assumption. Int J Epidemiol. 2017;46:1985-1998.

12. Koller M, Stahel WA. Sharpening Wald-type inference in robust regression for small samples. Computational Statistics & Data Analysis. 2011;55:2504-2515.

13. Burgess S, Bowden J, Dudbridge F, Thompson SG. Robust instrumental variable methods using multiple candidate instruments with application to Mendelian randomization. arXiv preprint arXiv:160603729. 2016

14. Zhao Q, Wang J, Bowden J, Small DS. Statistical inference in two-sample summary-data Mendelian randomization using robust adjusted profile score. arXiv preprint arXiv:180109652. 2018

15. Verbanck M, Chen C-Y, Neale B, Do R. Detection of widespread horizontal pleiotropy in causal relationships inferred from Mendelian randomization between complex traits and diseases. Nat Genet. 2018;50:693.

16. Qi G, Chatterjee N. Mendelian randomization analysis using mixture models for robust and

efficient estimation of causal effects. Nature Communications. 2019;10:1941.

17. Sinnott-Armstrong N, Tanigawa Y, Amar D *et al.* Genetics of 38 blood and urine biomarkers in the UK Biobank. BioRxiv. 2019660506.

18. Mahajan A, Taliun D, Thurner M *et al.* Fine-mapping type 2 diabetes loci to single-variant resolution using high-density imputation and islet-specific epigenome maps. Nat Genet. 2018;50:1505.
